# Evidence for the independent evolution of a rectal complex within the beetle superfamily Scarabaeoidea

**DOI:** 10.1101/2024.02.27.582323

**Authors:** Robin Beaven, Barry Denholm, Maria Fremlin, Davide Scaccini

## Abstract

Rectal or cryptonephridial complexes have evolved repeatedly in arthropods, including in beetles where they occur in ∼190,000 species of Cucujiformia and Bostrichoidea, and Lepidoptera where they occur in ∼160,000 species. Sections of the Malpighian/renal tubules coat the outer surface of the rectum, acting as powerful recycling systems of the gut contents, recovering water and specific solutes. There are hints that a rectal complex evolved independently within another beetle group, Scarabaeoidea. Here we report our observations of rectal complexes in Scarabaeoidea, which support this view. We did not find a rectal complex in the related group, Staphylinoidea, or in Lucanidae, a basal group of Scarabaeoidea. We did observe rectal complexes in *Melolontha melolontha* (Melolonthini), *Pachnoda marginata* and *Cetonia aurata* (Cetoniinae), consistent with previous reports from these groups. Intriguingly we found that rectal complexes occur in adult, but not *M. melolontha* larvae, and larvae but not adults within Cetoniinae, indicating dramatic pupal remodelling of these organ systems. Insights into the structure of the rectal complexes of Scarabaeoidea are compared with the well-studied rectal complexes of Cucujiformia. Finally we discuss possible functions of the rectal complexes of beetles within Scarabaeoidea, and future approaches to address this question.

## 1. Introduction

Cryptonephridial complexes (CNC), also known as rectal complexes, are remarkably widespread organ systems which appear to have evolved multiple times in arthropods. They result from an evolutionary reorganisation of the renal Malpighian tubules (MpTs) in relation to the rectum. MpTs function in maintaining water and solute homeostasis, and removal of nitrogenous waste (Denholm, 2013; Dow et al., 1994; Wessing and Eichelberg, 1978). Their ancestral organisation is as blind ended tubes, the proximal ends of which open into the gut at the junction of the midgut and hindgut, allowing the urine they produce to be expelled. In the derived, cryptonephridial organisation, their distal ends form a secondary association with the gut, having highly elaborated courses over the surface of the rectum, where they function in the retrieval of water or solutes from the rectal contents, to be returned to the haemolymph (Kolosov and O’Donnell, 2019; Ramsay, 1964). The CNC is ensheathed in a tissue, the perinephric membrane, which insulates the complex to enable its efficient function (Grimstone et al., 1968; Koefoed, 1971). There are also some cases of more rudimentary complexes which lack a perinephric membrane (Arab and Caetano, 2002; Dallai et al., 1991; Ullman et al., 1989). We will use rectal complex as a broader term, with CNC reserved for complexes with a perinephric membrane.

The largest group in which CNCs are found is in the beetle Cucujiformia infraorder and Bostrichoidea superfamily, for which there seems to be a single evolutionary origin. These groups contain ∼190,000 species (Hunt et al., 2007), likely to possess CNCs. The structure and physiology of the CNC has been long studied and extensively characterised in beetles from the family Tenebrionidae (infraorder Cucujiformia), particularly in *Tenebrio molitor* and *Onymacris* species (Coutchié and Machin, 1984; Dunbar and Winston, 1975; Grimstone et al., 1968; Hansen et al., 2004; Machin, 1975; Machin and O’Donnell, 1991; Marshall and Wright, 1972; Noble-Nesbitt, 1970; O’Donnell and Machin, 1991; Ramsay, 1964). More recently a molecular understanding of CNC function has been gained using the model organism *Tribolium castaneum*, which has the same CNC structure and physiology as other Tenebrionid beetles (Beaven et al., 2023; King and Denholm, 2014; Naseem et al., 2023). The current understanding is that potassium chloride is actively transported into the cryptonephridial MpTs from the haemolymph (Grimstone et al., 1968; Machin and O’Donnell, 1991; Naseem et al., 2023; Tupy and Machin, 1985). This is likely to occur via specialised cells, the leptophragmata, which are positioned beneath holes in the perinephric membrane, giving them access to the haemolymph (Grimstone et al., 1968; Lison, 1937; Naseem et al., 2023; Ramsay, 1964). This generates high osmolarity within the complex, drawing water out of the rectal contents (Machin, 1979; Ramsay, 1964). Fluid flows out of the cryptonephridial complex along the MpT lumen, into regions of the MpTs which lie freely within the body cavity, where reclaimed water can be returned to the haemolymph (Ramsay, 1964). The flow of contents within the rectum and within the cryptonephridial MpTs are in opposite directions, creating a countercurrent organisation to maximise water return (Kirschner, 1967; Phillips, 1970). The CNC has therefore enabled survival in arid environments (Beaven et al., 2023).

A remarkably similar system seems to have evolved independently within Lepidoptera, where it is present in the larval stage. CNCs have been observed in the clade Ditrysia (Henson, 1937; Irvine, 1969; Ito, 1921; Judy, 1968; Metalnikov, 1908; O’Donnell and Ruiz-Sanchez, 2015; Ramsay, 1976; Saini, 1964; Wigglesworth, 1974), which contains most species (∼160,000; Regier et al., 2009), and only seems to be absent in the most basal, species-poor groups (Shields et al., 2006). Interestingly, although CNCs can return water to the body in Lepidoptera (Reynolds and Bellward, 1989), it is considered that their main function is in the return of solutes from the rectal contents, in order to maintain correct acid-base balance. This could be an adaptation to allow rapid digestion of leaves during the larval stage (Kolosov and O’Donnell, 2019; Ramsay, 1976). Therefore CNCs appear to be powerful recycling systems, which have evolved to meet the diverse environmental and dietary challenges faced by different insect groups.

A rectal complex has also been reported in some beetle species within Scarabaeoidea, with the most detailed description being for adults of the common cockchafer, *Melolontha melolontha* (tribe Melolonthini; Fig. 1). In this species the rectal complex is reported to be ensheathed in a very fine cellular layer which may represent a perinephric membrane (Lison, 1938; Saini, 1964). The picture of the presence of rectal complexes across Scarabaeoidea remains extremely unclear, as differences in morphologies may reflect evolutionary changes between species, but also developmental changes between stages studied (larval or adult). Furthermore there has been no attempt to look systematically across this group, and the quality and detail of previous reports varies significantly. No rectal complex was observed in adults of related superfamilies Staphylinoidea or Hydrophiloidea (Dufour, 1824). Rectal complexes also appear to be lacking in several families and subfamilies within Scarabaeoidea (Fig. 1), namely in species of Lucanidae, in adults and larvae (Dufour, 1824; Dufour, 1842; Edwards, 1930; Saini, 1964), and in adults of Geotrupidae (Saini, 1964), Scarabaeinae (Dufour, 1842; Verma, 1969), and Aphodiinae (Verma, 1969).

**Figure 1.**
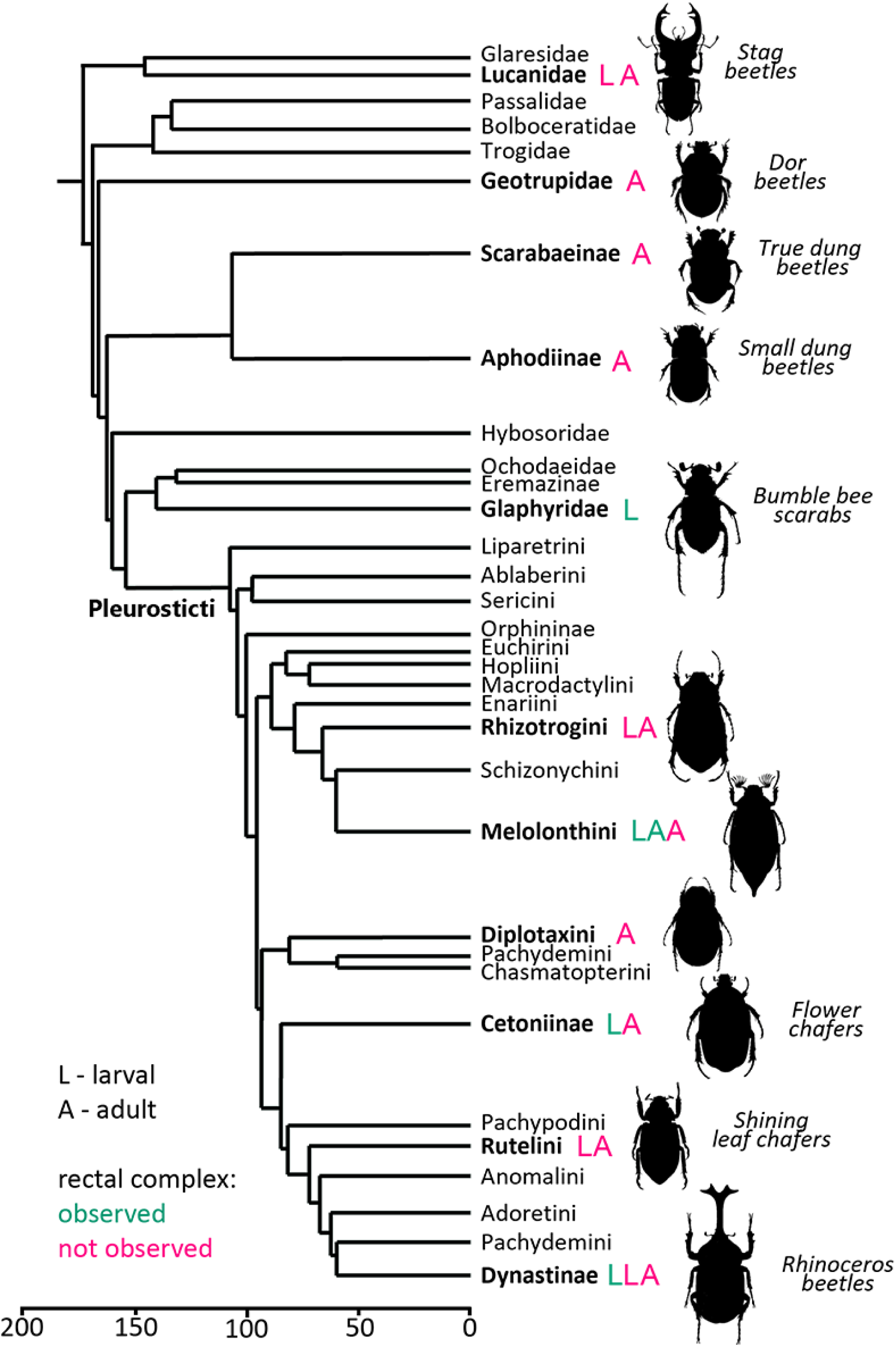
Phylogenetic tree of the beetle superfamily Scarabaeoidea, with previous reports of occurrence of rectal complex. The tree is drawn according to Ahrens et al., 2014, and represents an ∼180 million year evolutionary period. Reports of CNC or rectal complex presence (green) or absence (magenta) are indicated, along with the life stage reported (L – larval stage, A – adult stage). Note that parts of Pachydemini appear in two different branches. Scale bar shows estimated evolutionary time in millions of years.

Here we report our initial observations of gut and MpT morphology in beetle species within Scarabaeoidea, from which we are beginning to tackle these questions. The phylogenetic distribution we observed, along with significant morphological differences in comparison to rectal complexes observed in Cucujiformia and Bostrichoidea, support the notion that a rectal complex evolved independently within Scarabaeoidea. Our findings shed further light on the morphology of this complex in Scarabaeoidea, particularly the question of the presence or absence of a perinephric membrane. Furthermore, interestingly morphological differences were seen between larvae and adults, indicating rectal complex remodelling during pupal metamorphosis. This study provides a useful starting point for more extensive investigations. These will extend our understanding of the evolution of rectal complexes, and their contribution to the evolutionary success of the insect groups in which they occur, as well as providing a greater understanding of the adaptations of Scarabaeoidea, which is a very large and ecologically important beetle superfamily.

## 2. Materials and Methods

*Platycerus caraboides* larvae were directly collected from deadwood of deciduous trees in the Vicenza province of Italy (Scaccini, 2016; Scaccini, 2022). A *Dorcus parallelipipedus* adult and *M. melolontha* larva (third instar) were collected in Colchester, UK, with the *D. parallelipipedus* being from a population we have previously studied (Scaccini et al., 2023). The *M. melolontha* adult was collected from Shottisham, Suffolk, UK. *Nicrophorus vespilloides* larvae were from a lab population, established from wild caught individuals from Lincoln, UK. *Pachnoda marginata* were obtained commercially as larvae from Livefood UK Ltd. *Cetonia aurata* larvae and adult females were collected in Colchester, UK from a previously described population (Fremlin, 2018), with both being used, as well as their offspring. In the case of *C. aurata* and *P. marginata*, larvae were kept in leaf mould with added rotten wood, both from deciduous trees, in plastic boxes with drilled airholes, at 25°C. The substrate was kept moist by spraying with water every few days. Adults were kept in the same conditions and fed with chopped apple and banana.

Live beetles were decapitated with scissors, and where possible the central nervous system removed with forceps. They were then dissected in PBS and imaged on a Leica S9i microscope with integrated camera. Fixation of the dissected *M. melolontha* gut and MpTs was in 4% formaldehyde in PBS at 4°C overnight, before washing in PBT+BSA (PBS + 0.3% Triton X-100 + 0.5% BSA). Staining was with anti-α-Tubulin (mouse, 1:20, AA4.3, DSHB) for 4 hrs at room temperature, washing in PBT+BSA followed by staining with anti-mouse-488 (1:200, Jackson ImmunoResearch), phalloidin-568 (1:200, A12380, Molecular Probes) and DAPI (1:1000, Molecular Probes) for 5 hrs at room temperature. The sample was placed on a coverslip, covered in mounting media (85% glycerol, 2.5% propyl gallate), and imaged on a Nikon A1R confocal microscope.

## 3. Results

### 3.1. Rectal complexes are not observed in Staphylinoidea and basal Scarabaeoidea

The burying beetle, *Nicrophorus vespilloides*, is a member of the superfamily Staphylinoidea which is the sister group to Scarabaeoidea (Fig. S1), and would not be expected to have a rectal complex, assuming the rectal complex evolved independently within Scarabaeoidea. This is indicated by reports from adult *Staphylinus erythropterus*, *Paederus riparius*, and *Silpha obscura* (Dufour, 1824), which are members of Staphylinoidea. For final instar larvae of *N. vespilloides* we found that a looped midgut sits ventral to the hindgut (Fig. 2A). The MpTs are highly elaborated and lie alongside the hindgut, but are not applied to its surface (Fig. 2B,C). This supports the lack of a rectal complex in Staphylinoidea, and is the first observation we are aware of in the larval stage. The MpTs insert laterally into each side of the gut (Fig. 2D). For this species and for all species subsequently reported, the point of insertion is assumed to be the junction of the midgut and hindgut, which is consistently the case across insects. Two MpTs were observed (Fig. 2C,D). In Coleoptera the number of MpTs generally ranges from four to eight (Beutel and Haas, 2000). Staphylinoidea have mostly been considered to have four (Lawrence and Newton, 1982), although members of the tribe Nanosellini were reported to have only two. This was speculated to represent a reduced number due to miniaturisation of body size in this group (Polilov, 2008), but this question should be revisited in light of our observation in a larger species.

**Figure 2.**
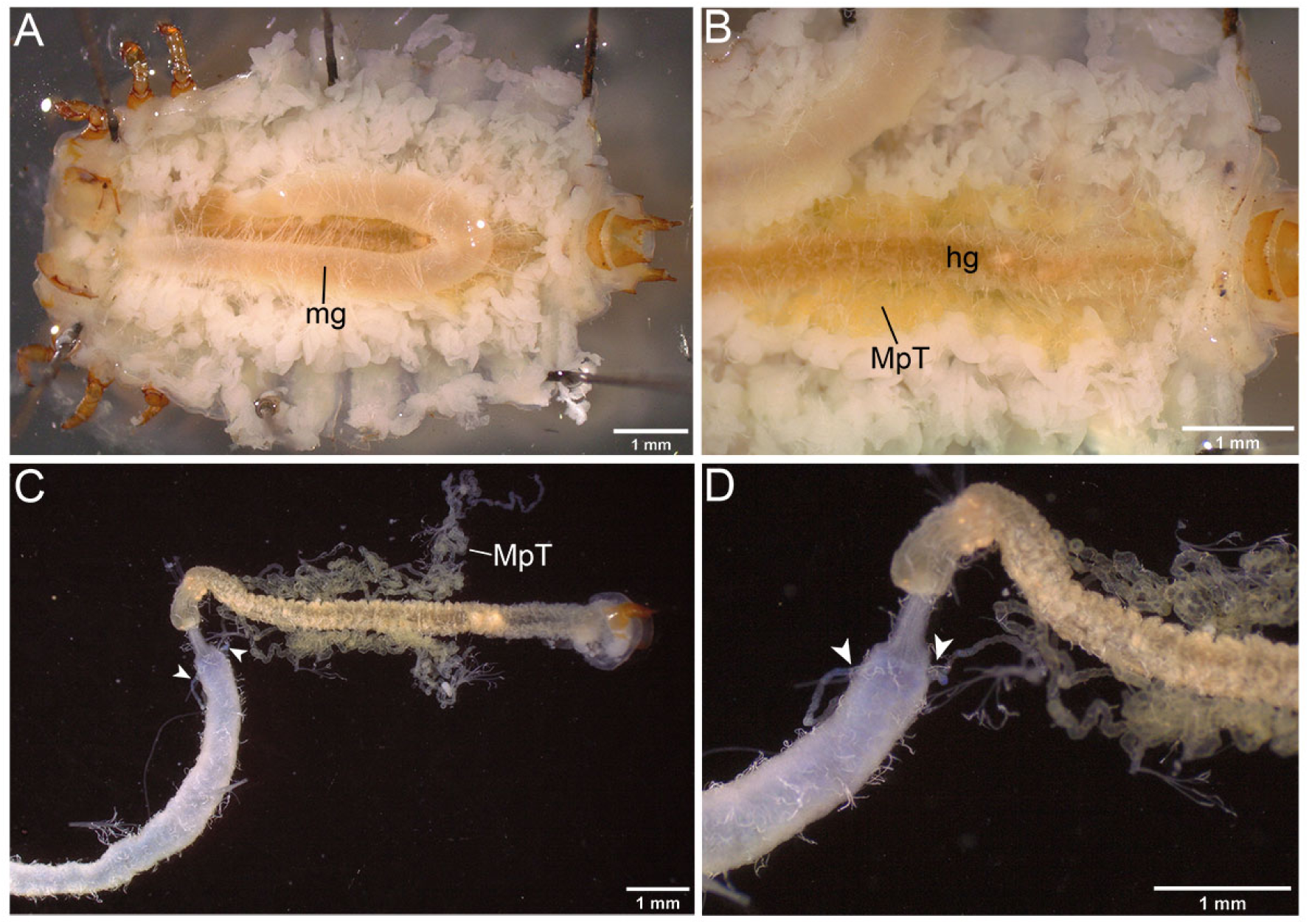
Gut and Malpighian tubules of final instar larva of the burying beetle, *Nicrophorus vespilloides*. In all images anterior is to the left. **(A)** Ventral view in which the looped midgut (mg) can be seen lying on top of the hindgut. **(B)** Ventral view in which the midgut has been lifted to reveal the hindgut (hg), and the mass of MpTs (MpT) lying on either side of it. **(C)** Gut removed from body, revealing two MpTs lying on either side of the hindgut, with insertion points into the gut indicated (arrowheads). **(D)** An enlarged view with MpT insertion into the gut marked (arrowheads).

Lucanidae, the stag beetles, is a basal family within Scarabaeoidea (Fig.1). Reports of adults of the European stag beetle, *Lucanus cervus* (Dufour, 1824; Saini, 1964), and larvae and adults of the lesser stag beetle, *Dorcus parallelipipedus* (Dufour, 1842; Edwards, 1930; Saini, 1964), suggest that members of this family lack a rectal complex. In final (3^rd^) instar larvae of another stag beetle species, *Platycerus caraboides*, we found caeca in three regions of the gut (Fig. 3A,B). There are some caeca like extensions at the anterior end of the midgut (Fig. 3B). There is also a central row of caeca within the midgut and posterior caeca within the hindgut (Fig. 3A,B). MpTs wrap around the ventral caeca from the posterior row (Fig. 3C). There are four MpTs, with two inserting into the gut on the dorsal side (Fig. 3D), and two on the ventral side, which was revealed when the MpT regions enwrapping the caeca were teased away (Fig. 3E). Highly elaborated MpTs coat the surface of part of the midgut on the dorsal side (Fig. 3F). The most anterior part of the hindgut is broad, and is followed posteriorly by a narrow ileum. Posterior to this is a dilated colon (Fig. 3A), likely to be a smaller equivalent of the fermentation chamber which exists in larvae of the family Scarabaeidae (Chapman, 1998). MpTs are associated with the ileum and colon, although this does not appear to represent a rectal complex, as MpTs are only loosely associated and are not extensively elaborated (Fig. 3G). The observed morphology looks broadly comparable to that reported for larval *D. parallelipipedus* by Edwards (in which the posterior caeca are referred to as diverticulum of the ileum) (Edwards, 1930).

**Figure 3.**
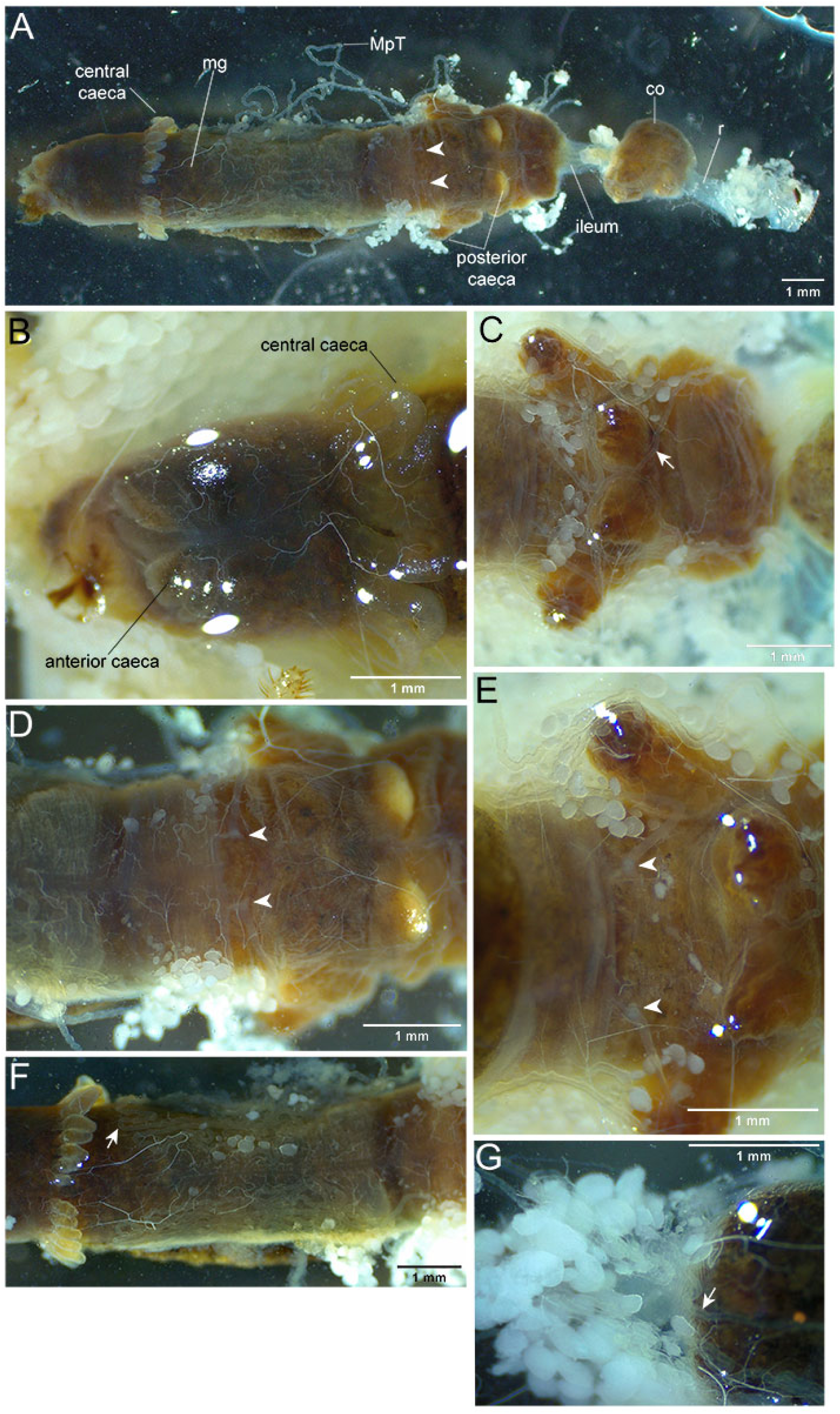
Gut and Malpighian tubules of final instar larvae of a stag beetle, *Platycerus caraboides*. In all images anterior is to the left. **(A)** Dorsal view. Overview of the dissected gut with midgut (mg), ileum, colon, rectum (r), central and posterior rows of caeca, and insertion point of two of the MpTs (arrowheads) indicated. **(B)** Ventral view. Anterior and central caeca are indicated. **(C)** Ventral view. MpTs enwrap posterior caeca on ventral side of gut (arrow). **(D)** Dorsal view. Two MpTs insert into dorsal side of gut (arrowheads). **(E)** Ventral view. Two MpTs insert into ventral side of gut (arrowheads). **(F)** Dorsal view. Highly elaborated MpTs cover part of the dorsal midgut (arrow). **(G)** Ventral view. Elaborated MpTs (arrow) loosely associate with the ileum and colon.

We also dissected an adult female lesser stag beetle (*D. parallelipipedus*), to compare with previous reports. We observed elaborated MpTs loosely surrounding the midgut and hindgut (Fig. 4A,B). Four MpTs insert into the gut (Fig. 4C). There is only a very loose association between the MpTs and the rectum (Fig. 4B). Together with our findings from *P. caraboides*, this supports previous suggestions that a rectal complex is not present in species within Lucanidae. The absence of a rectal complex in Staphylinoidea, as well as in Lucanidae (which is a basal family of Scarabaeoidea), support the view that the rectal complex of Scarabaeoidea has an independent evolutionary origin from the CNC of Cucujiformia and Bostrichoidea (see Fig. 1 and Fig. S1).

**Figure 4.**
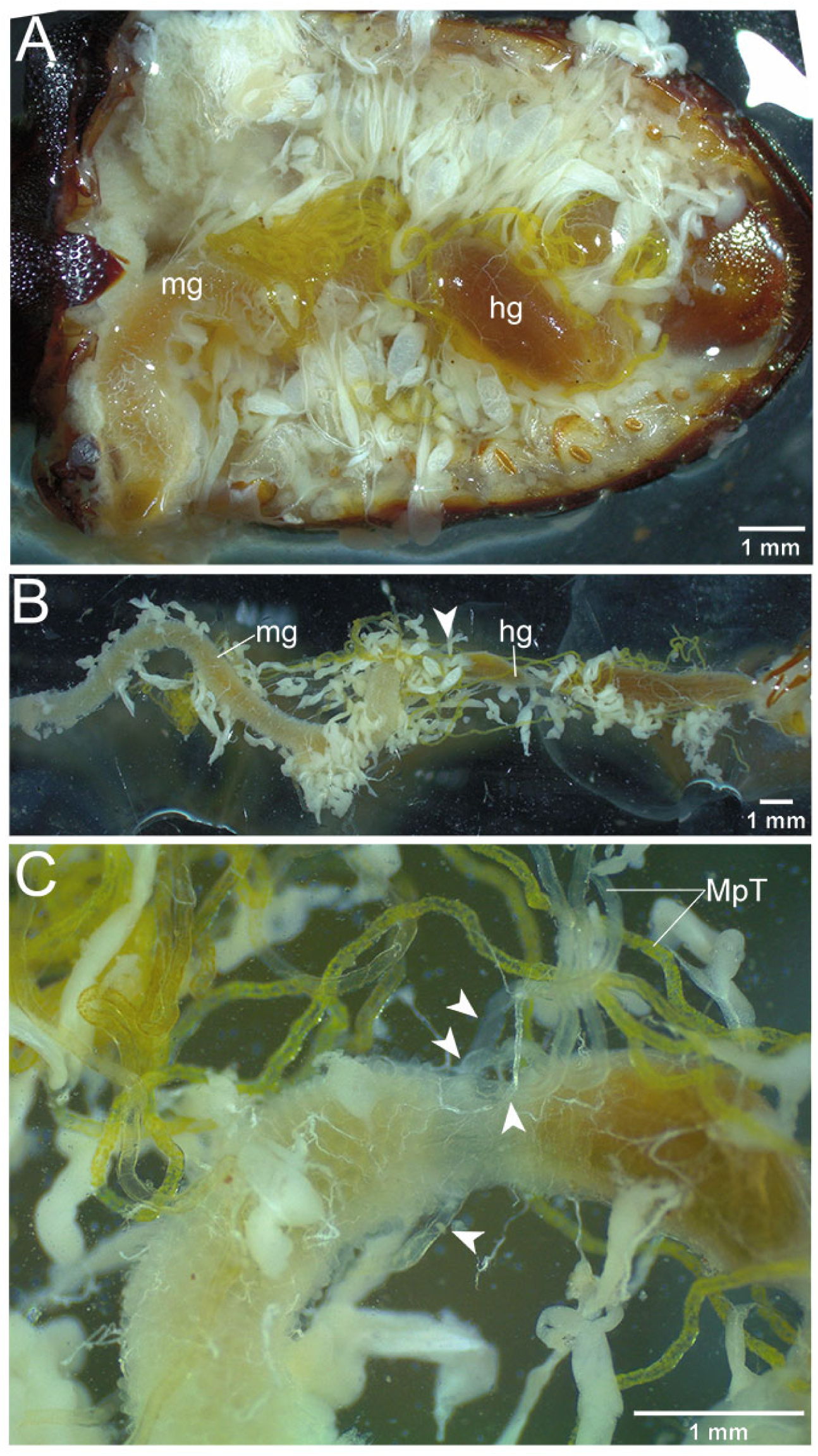
Gut and Malpighian tubules of an adult lesser stag beetle, *Dorcus parallelipipedus*. In all images anterior is to the left. **(A)** Dorsal view indicating position in the body cavity of midgut (mg) and hindgut (hg). **(B)** Dissected gut and associated MpTs (which have yellow and transparent regions), with midgut and hindgut indicated as in A, along with the position of the midgut-hindgut boundary (arrowhead). **(C)** Close up view of the midgut-hindgut boundary, with the four MpTs indicated (arrowheads) close to their point of entry into the gut.

### 3.2. The rectal complex of *Melolontha melolontha* likely forms during pupal metamorphosis

It was previously reported that adult cockchafers, *Melolontha melolontha*, have a rectal complex. This is surrounded by a fine cellular layer, which may represent a rudimentary perinephric membrane (Lison, 1938; Saini, 1964). The morphology of the larval gut and MpTs has not been reported before. In a *M. melolontha* larva, we observed two morphologically distinct rows of caeca in the midgut (Fig. 5A-C). The hindgut is folded into an S-shape. There is a very narrow ileum (i.e. the anterior portion of the hindgut), which opens into a huge fermentation chamber (Fig. 5C). The fermentation chamber is folded back to lie dorsally upon the ileum (Fig. 5A,B). As reported for other beetles from Scarabaeoidea (Kucuk et al., 2023; Sheehan et al., 1982), the bases of papillae can be seen on the surface of the fermentation chamber (Fig. 5A,B). The rectum can be observed as a fairly narrow section of the most posterior hindgut, forming two lobes, and it runs over the surface of the fermentation chamber to the anus (Fig. 5A,B). There are elaborated MpTs over some of the surface of the midgut (Fig. 5D). There are also some elaborated MpTs sandwiched between the rectum and fermentation chamber, revealed when lifting the rectum away from the fermentation chamber (Fig. 5E), however this does not appear to represent a rectal complex, as there is a relatively small amount of MpT, and this has very limited coverage of the rectal surface. Two MpTs insert into the gut side by side on its ventral surface, close to the posterior row of caeca (Fig 5F). Two more MpTs insert dorso-laterally on either side of the gut (Fig. 5G). Curiously the dorsal pair lie some way from the row of caeca. It appears that all four MpTs insert at the boundary between lighter and darker gut regions, likely to be the midgut and hindgut respectively. This boundary is not straight, forming a chevron on the ventral side of the gut, meaning the row of caeca does not follow the midgut-hindgut boundary (Fig. 5F,G).

**Figure 5.**
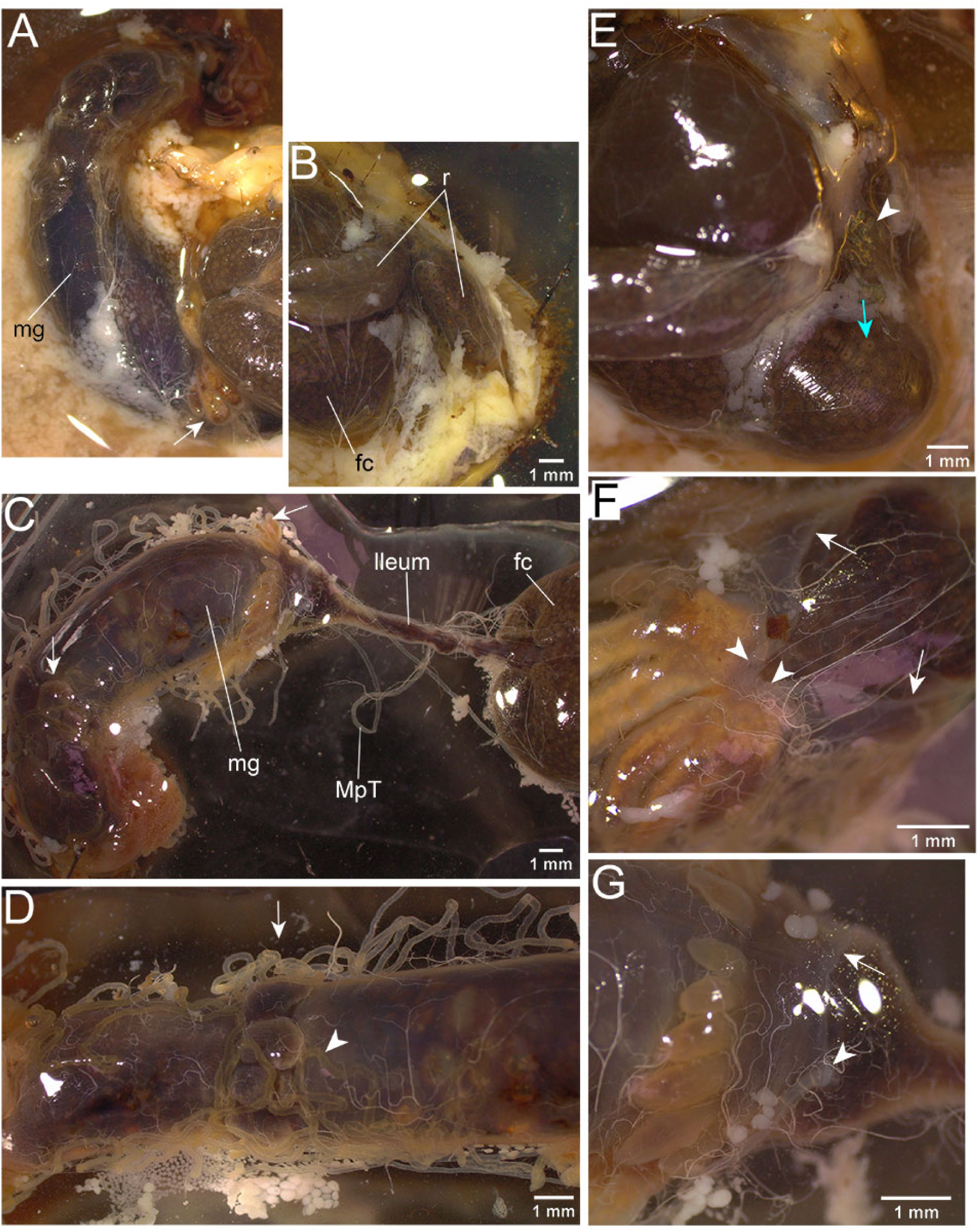
Gut and Malpighian tubules of a final instar larva of the cockchafer, *Melolontha melolontha*. In all images anterior is to the left. **(A,B)** Panels together forming an overview of the gut. Ventral view with midgut (mg), hindgut fermentation chamber (fc) and two lobes of rectum (r) indicated, along with the posterior row of caeca (white arrow) which is partially beneath the fermentation chamber. **(C)** Dorsal view in which the hindgut has been unfolded, with the midgut (mg), ileum, fermentation chamber (fc) and two rows of caeca (white arrows) indicated. The posterior row of caeca are close to the midgut-hindgut junction. **(D)** Dorsal view of a midgut portion, showing the anterior row of caeca (arrow), and convoluted MpTs associated with the midgut (arrowhead). **(E)** Ventral view with most posterior lobe of rectum lifted, revealing a region of elaborated MpTs (arrowhead) which had been sandwiched between the rectum and the fermentation chamber. The bases of papillae dapple the fermentation chamber surface (blue arrow). **(F)** Ventral view showing junction of two MpTs with the gut (arrowheads), and the boundary between the lighter midgut and darker hindgut (arrows). **(G)** Dorso-lateral view showing the insertion into the gut of one of the lateral pair of MpTs (arrowhead) and the midgut-hindgut boundary (arrow).

Regarding the rectal complex, our observations were unexpected, as while there are clearly elaborated MpTs associating closely with the hindgut (Fig. 5E), this does not resemble a rectal complex, although the presence of a rectal complex has been reported in adults. Even though there are indications that rectal complexes may be present in larvae of other species within Melolonthini, the detail in these reports is not sufficient to make clear comparisons with our findings for *M. melolontha*, and further studies would be required to clarify the occurrence of rectal complexes in larvae of Melolonthini.

There are also white regions of MpT (black arrows). Part of male reproductive organs (*) are also indicated. **(B)** The rectal complex. **(C)** Enlarged region corresponding to blue boxed rectangle in B. A thick white MpT (black arrow) and a narrower clear MpT (black arrowhead) are indicated. **(D)** Anterior section of midgut, along the surface of which run branched MpTs (blue arrow). **(E)** Central region of hindgut, with associated branched MpTs (blue arrow). **(F)** The four MpTs indicated (white arrowheads) close to their insertion points into the gut. This image is from fixed tissue, which has altered its colouration.

To confirm the apparent difference between larvae and adults, we dissected an adult male *M. melolontha*. This showed a dramatically remodelled gut and MpTs, when compared to the larva of the same species. Convolutions in the gut bring a section of the hindgut into close proximity to the rectum (Fig. 6A). Three morphologically distinct MpT regions were observed. There are pale yellow MpTs with a branched morphology, and some of these are sandwiched between the rectum and the loop of the hindgut (Fig. 6A). There are also unbranched MpT regions, which are either white, and of a variable and generally wider diameter, or colourless and of a generally narrower diameter (Fig. 6A-C). These changes to MpT morphology between larvae and adults have been reported for the related species *P. anxia*. In this species the larval MpTs are morphologically uniform but in the adult transform to a morphology with three distinct regions, including a branched region referred to as diverticular (Berberet and Helms, 1972).

**Figure 6.**
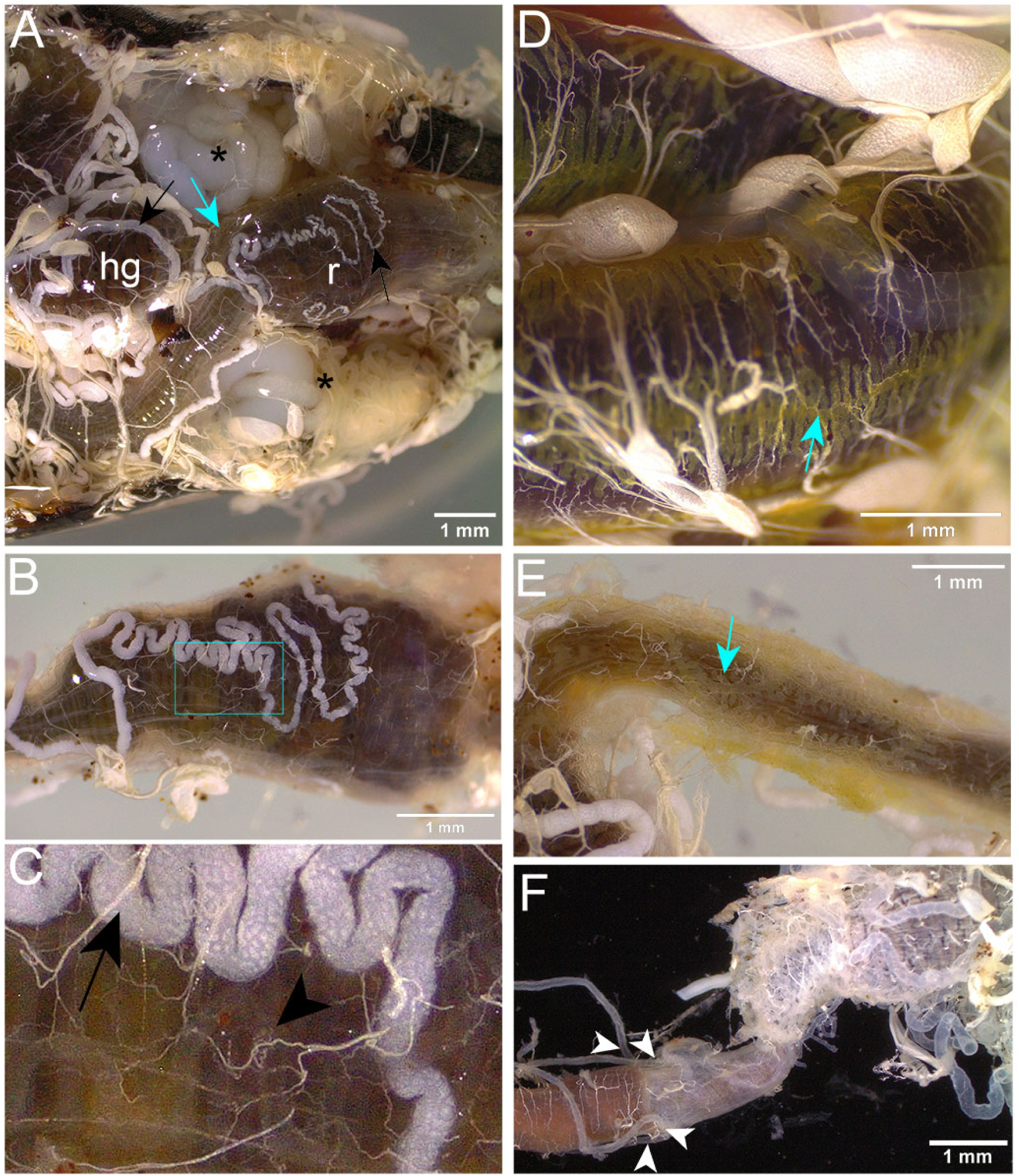
Gut and Malpighian tubules of an adult male cockchafer, *Melolontha melolontha*. All images are dorsal view, with anterior to the left. **(A)** Part of the hindgut (hg) is looped back into close proximity with the rectum (r) and highly branched MpTs are packed between the two (blue arrow).

We observed that both white and colourless MpTs are incorporated into a rectal complex, with elaborate sinuous courses over the surface of the rectum (Fig. 6A-C). Much of the gut has branched MpTs running along its length, intimately associated with its surface. This includes both midgut (Fig. 6D) and hindgut (Fig. 6E) regions. The branched MpTs have short regularly spaced branches along their length, which often arranged into opposite pairs (Fig. 6D,E). There are four MpTs which insert separately into the gut (Fig. 6F). For an overview of the gut and associated MpT organisation, see Fig. S2.

To gain a more detailed view of the rectal complex of adult *M. melolontha*, we fixed and fluorescently stained the gut and MpTs. This revealed a number of interesting features of rectal complex structure. The overall organisation is as follows. Bands of circular muscle encircle the epithelial layer of the rectum (Fig. 7A,B). Overlaying this are the sinuous thick and thin MpT subtypes, which are both coated in nets of muscle (Fig. 7A,B). The branched MpT regions, lying outside the rectal complex, are also netted by muscle cells (Fig. 7C). There appear to be a number of other associated cell types. There is a cellular layer surrounding the rectal complex (Fig. 7A,B), which was just discernible as an extremely thin transparent membrane, in the unstained preparation. The membrane looks structurally very different from the perinephric membrane tissue observed in the Tenebrionid beetles, *Tribolium castaneum* and *Tenebrio molitor*. It does not appear to form a continuous sheet coating the complex (Fig. 7A,B), and would therefore not be expected to act as an insulating barrier. Trachea could be seen extending into the membrane (Fig. 7B), and the membrane and trachea may be part of a common tissue. Additionally other muscle cells were noted, which form connections with the thin MpTs and with the membrane tissue (Fig. 7A,B).

**Figure 7.**
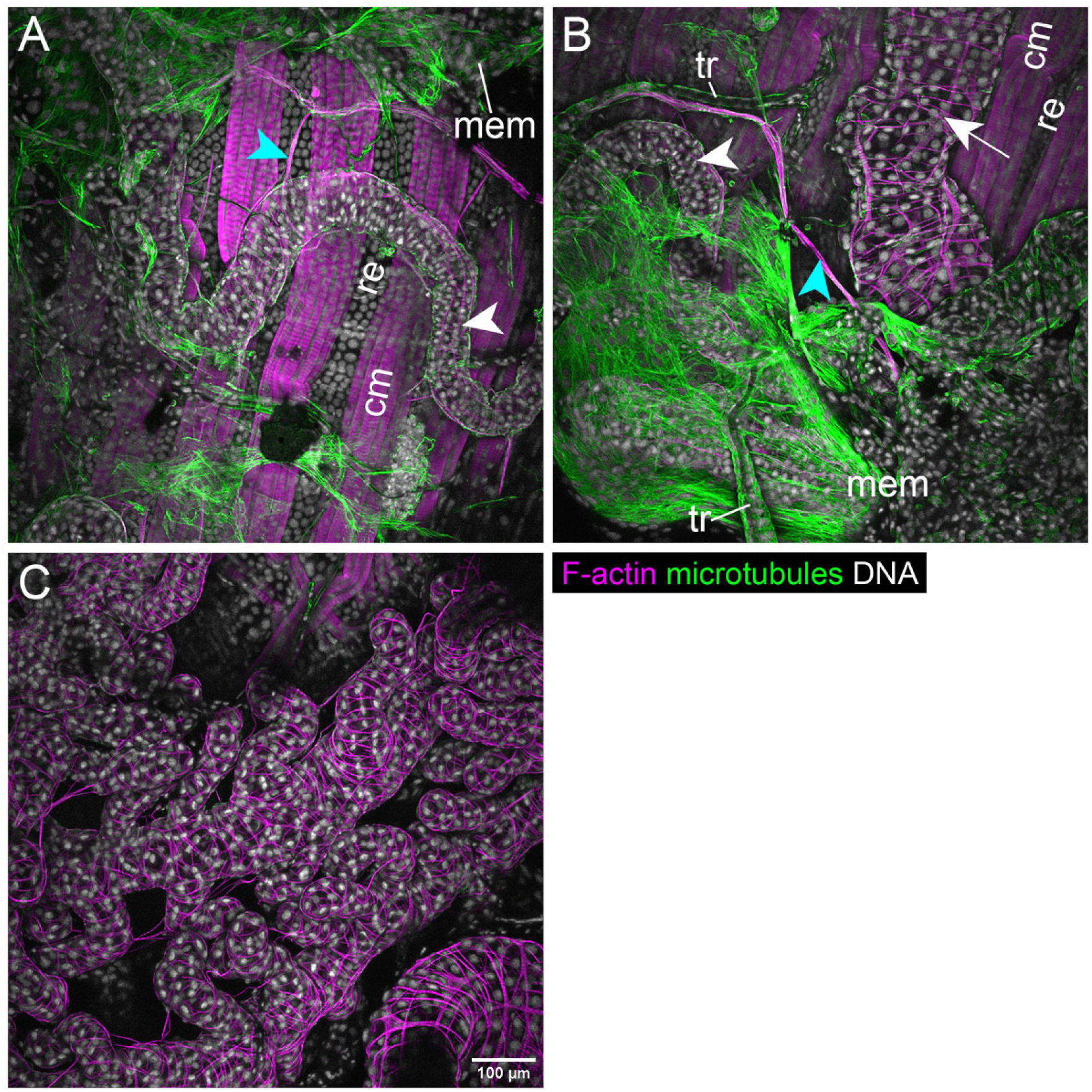
Fluorescent images of the rectal complex and branched MpTs from an adult male cockchafer, *Melolontha melolontha*. F-actin (phalloidin stain, magenta), microtubules (anti-alpha-Tubulin, green), and nuclei/DNA (DAPI stain, white). **(A)** The rectal epithelium (re) is encircled in bands of circular muscle (cm) around the circumference of the rectum. F-actin enrichment makes the muscle clearly visible. The thin MpTs (white arrowhead) are coated in a mesh of muscle, and have a sinuous course over the rectum. The complex is enwrapped by a cellular membrane layer (mem) although this does not appear to be a continuous sheet. This cellular layer is rich in microtubules. There are also more muscles (blue arrowhead), that form connections with the membrane tissue and the thin MpTs. **(B)** Labelled as in A, but also showing a region of a thick MpT (white arrow), and parts of the tracheal network (tr) which appear to connect to, or be part of, the membrane tissue. **(C)** Region showing the branched MpTs, netted in muscle.

### 3.3. A rectal complex is integrated into the hindgut fermentation complex of Cetoniinae larvae

We then investigated the morphology of the gut and MpTs in Cetoniinae, another subfamily within Scarabaeoidea, for which there were previous reports of a rectal complex in larvae but not adults (Dufour, 1824; Dufour, 1842; Werner, 1926). Firstly we investigated intestinal morphology in the rose chafer, *Cetonia aurata*. In a final (3^rd^) instar larva we observed that there are three rows of caeca within the midgut (Fig. 8A,B). Much like for *M. melolontha* larvae, the hindgut has an S-shaped course, with the rectum running ventrally over the surface of the fermentation chamber (Fig. 8A,B). However, unlike *M. melolontha*, the most posterior part of the rectum is enclosed by infoldings of the fermentation chamber from either side (Fig. 8B). A membranous tissue appears to overlay the seam made by the folded fermentation chamber. By pulling apart each side of the folded fermentation chamber, and lifting up the rectum, a remarkable structure was revealed. There are four rows of sinuous, tightly packed MpTs, which had entirely surrounded the rectum, and had been sandwiched between the rectum and fermentation chamber (Fig. 8C and Fig. 9A-C). Two pairs of MpTs enter this structure, one pair on either side of the rectum at its anterior end (Fig. 8C).

**Figure 8.**
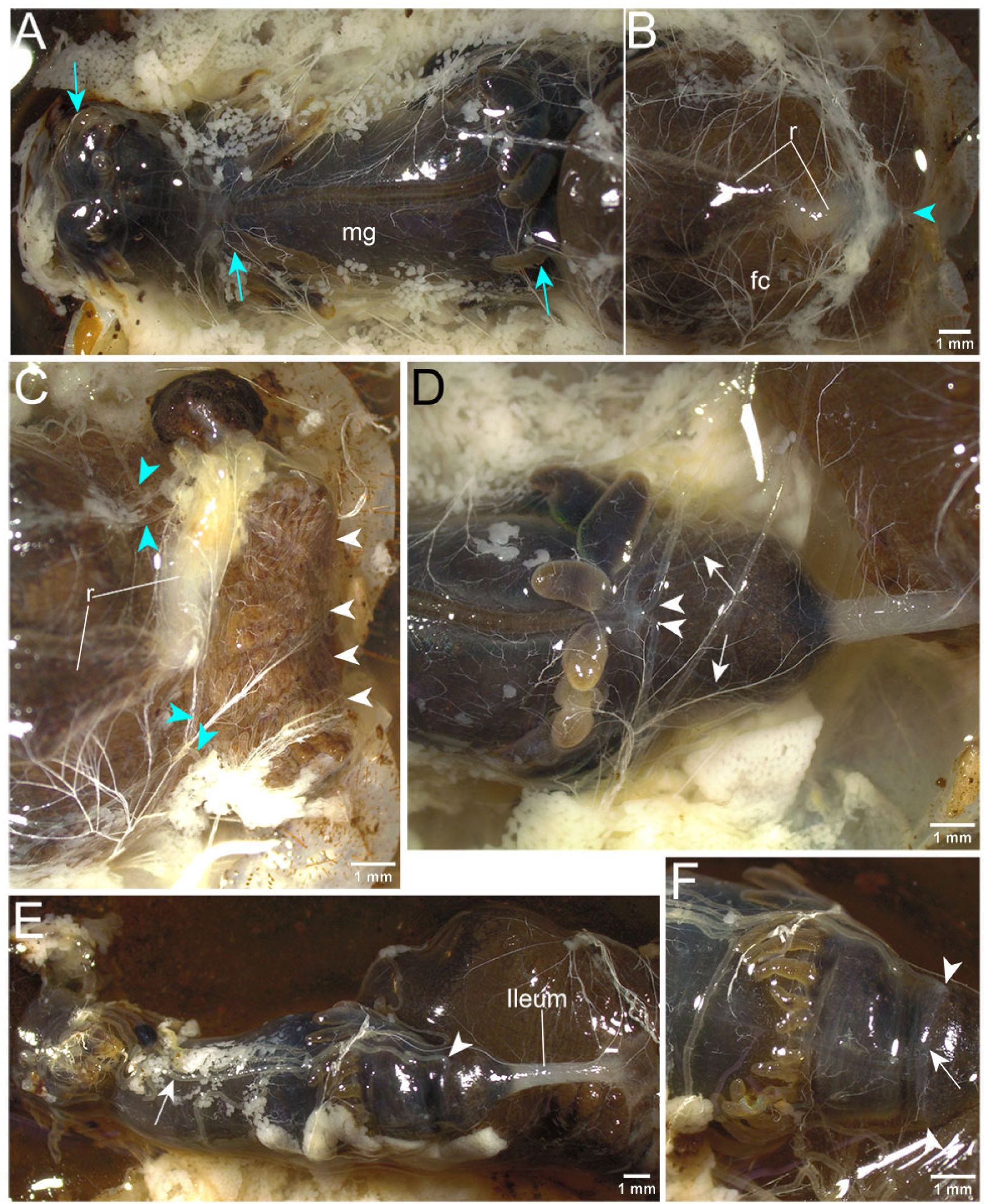
Gut and Malpighian tubules of a third instar larva of the rose chafer, *Cetonia aurata*. In all images anterior is to the left. **(A,B)** Panels together forming an overview of the gut. Ventral view, showing three rows of caeca (blue arrows) in the midgut (mg), as well as the fermentation chamber (fc), rectum (r) and the seam between the two infoldings of the fermentation chamber (blue arrowhead). **(C)** Ventral view. Upon opening the fold in the fermentation chamber and pulling aside part of the rectum (r), the sinuous course of four MpTs (white arrowheads) can be seen. A pair of MpTs enter on either side of the rectum (blue arrowheads). **(D)** Ventral view. Moving aside the fermentation chamber reveals the insertion points of two of the MpTs (white arrowheads), and the midgut-hindgut boundary (white arrows). **(E)** Dorsal view, showing ileum, one of the laterally inserted pair of MpTs (arrowhead) and a MpT associated with the midgut (arrow). **(F)** Dorsal view, showing insertion points of lateral pair of MpTs (arrowheads), at the midgut-hindgut boundary (arrow).

**Figure 9.**
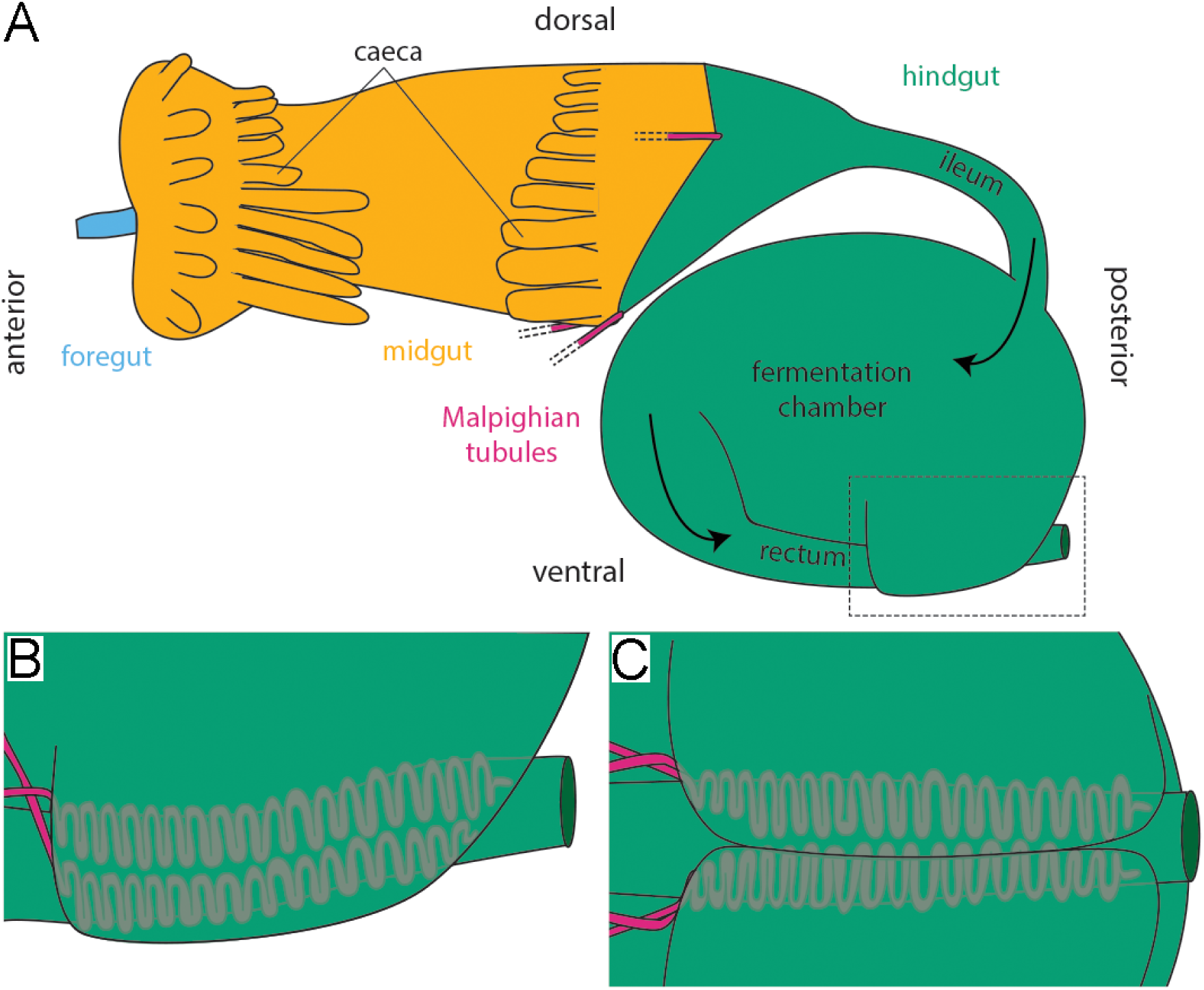
Approximate morphology of the gut and Malpighian tubules observed in Cetoniinae larvae. **(A)** Lateral view, with insertion points of MpTs shown. Arrows indicate the direction of flow of the gut contents. **(B)** Lateral view, showing incorporation of distal segments of MpTs into a rectal complex enclosed by infoldings of the fermentation chamber. Corresponds to boxed region in A. **(C)** Ventral view corresponding to boxed region in A.

The morphology and arrangement of the MpTs around the rectum looks strikingly similar to in the CNC of tenebrionid beetles (Grimstone et al., 1968; King and Denholm, 2014; Naseem et al., 2023; Ramsay, 1964), whilst its enclosure by the fermentation chamber is very different from what has been observed in other beetle groups. There also appears to be a transparent membrane structure, however this is sandwiched between the MpTs and the rectum, unlike the membrane seen in *M. melolontha* which resides outside the MpTs, as does the perinephric membrane seen in species of Tenebrionidae. As in *M. melolontha*, the anterior hindgut is a narrow ileum (Fig. 8D,E). The shape of the midgut-hindgut boundary, its position relative to the posterior row of caeca, and the insertion points of the four MpTs, also seems very similar between these groups (Fig. 8D-F). There are also some MpTs which associate with the midgut surface (Fig. 8E).

Dissecting adult *C. aurata* revealed dramatic remodelling of the gut and MpTs between larval and adult stages, as seen for *M. melolontha*. There are distinct MpT regions, with most being clear or pale yellow, along with a second type which are white (Fig. 10A-C). Some MpT regions are free within the body cavity, whilst others are loosely associated with the gut, particularly a central portion of the hindgut (Fig. 10A-F). Whilst there is clearly association of MpTs with the rectum, these do not have the regular sinuous course, or tight association typical of a rectal complex (Fig. 10C,D). The insertion points of four MpTs are as a pair close together on the ventral gut surface, and another pair on either side of the gut (Fig. 10G).

**Figure 10.**
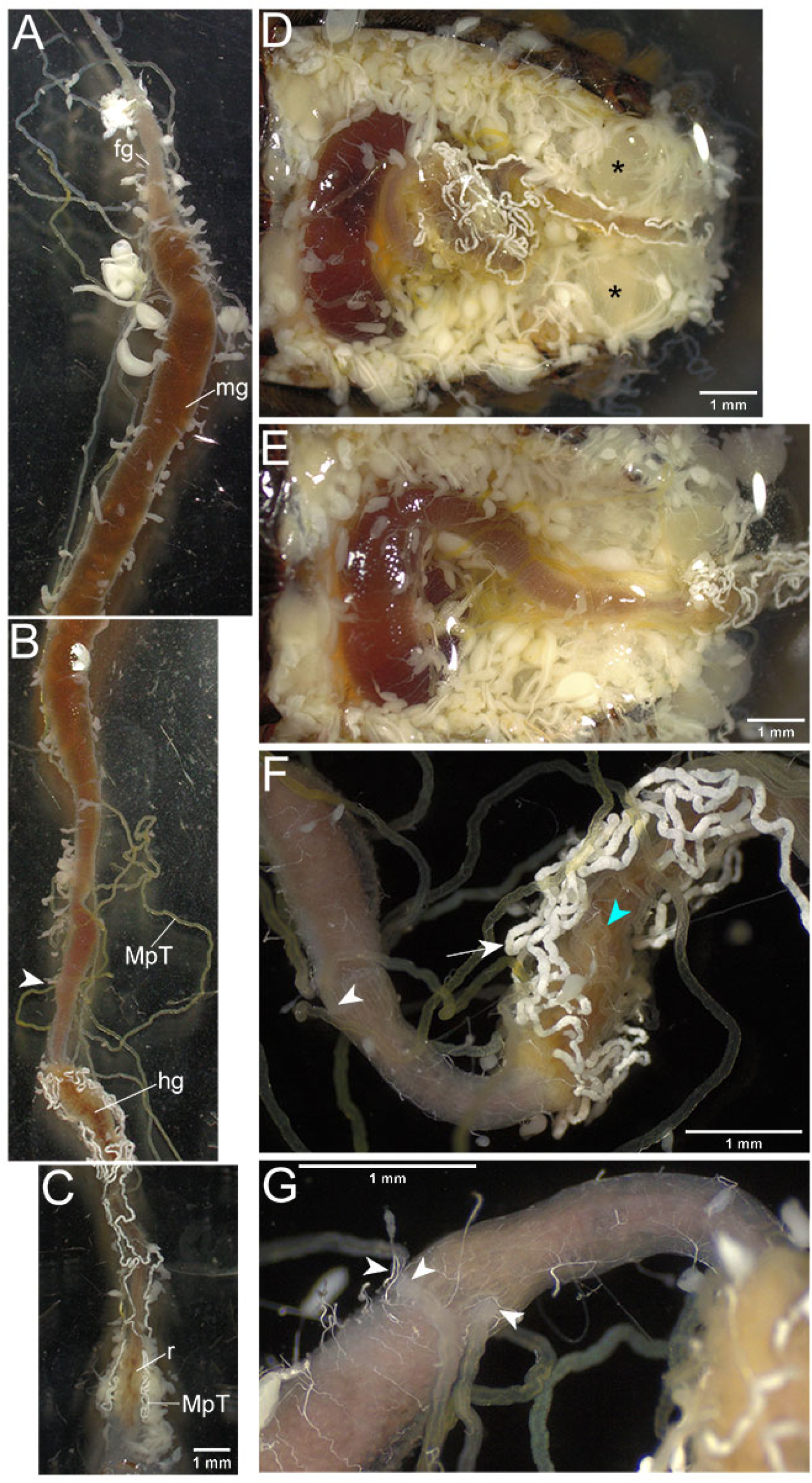
Gut and Malpighian tubules of an adult rose chafer, *Cetonia aurata*. **(A-C)** Dorsal views, anterior at the top, with foregut (fg), midgut (mg), hindgut (hg) and rectum (r) indicated. **(D)** Dorsal view to show position of gut within the abdomen. Part of male reproductive organs are indicated (*). **(E)** Dorsal view in which the hindgut has been unfolded to reveal the ileum. **(F)** Dorsal view, showing a region of the hindgut with elaborated, loosely associated MpTs. These are of a white subtype (white arrow) or transparent/yellow subtype (blue arrowhead). The insertion of one of the MpTs is indicated (white arrowhead). **(G)** Ventro-lateral view, showing a pair of MpTs which insert close together on the ventral side of the gut, and one of the laterally inserted pair of MpTs (arrowheads). Anterior is to the left in D-G.

Finally we investigated the gut and MpT morphology in *Pachnoda marginata*, which is another member of the subtribe Cetoniina, within the Cetoniinae subfamily. Dissecting final (3^rd^) instar larvae revealed a broadly similar morphology to *C. aurata* (Fig. 11A-F). The morphology at the adult stage also looked very similar between the two species (Fig. 12A-G), although in *P. marginata* there is a more extensive association of MpTs with the anterior part of the midgut (Fig. 12A). These findings show that an unusual rectal complex, integrated into the fermentation chamber, is more widespread in the subtribe Cetoniina. Investigation of further groups within and beyond the subfamily Cetoniinae will be required to reveal how widespread and variable this structure is.

**Figure 11.**
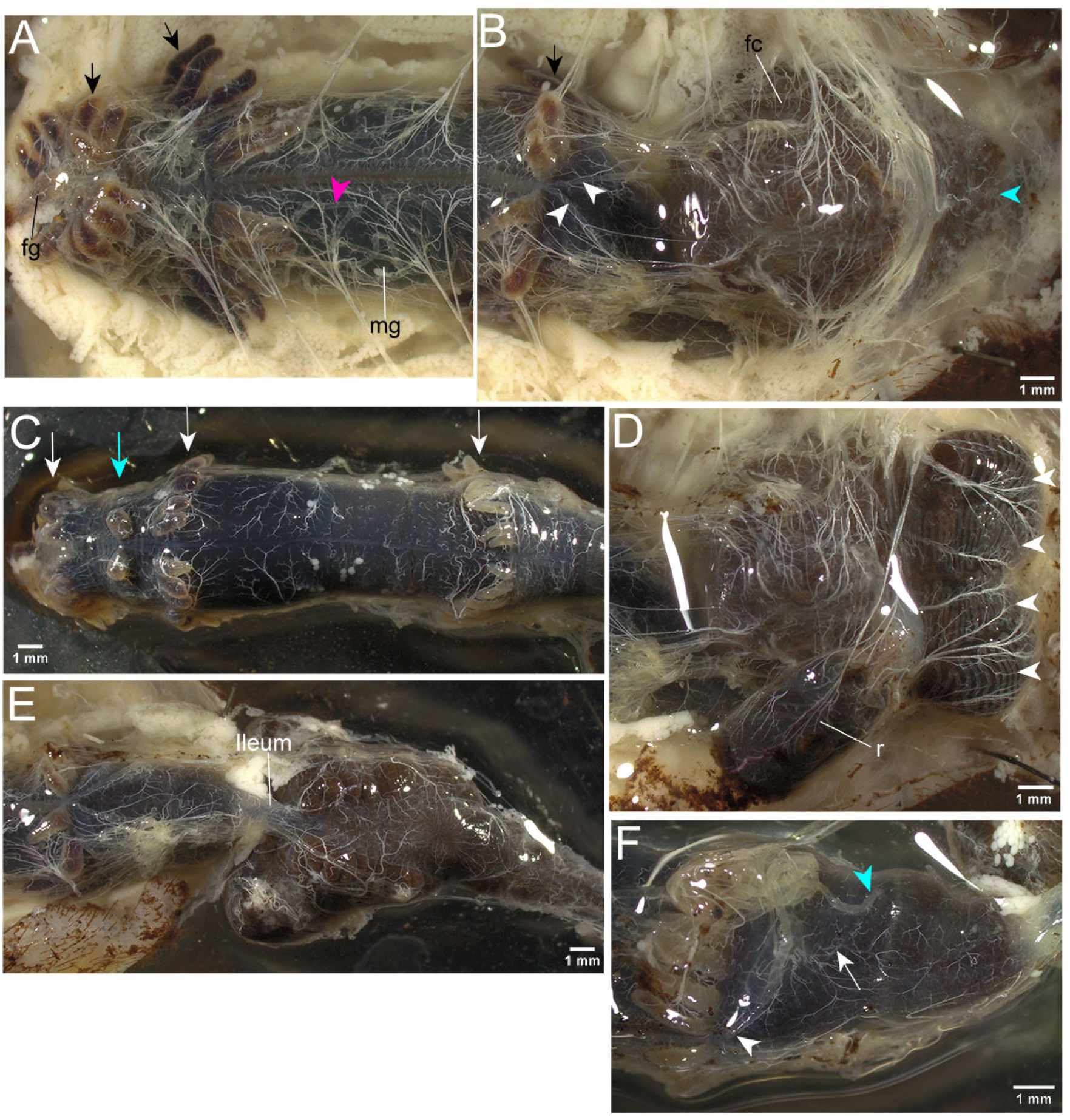
Gut and Malpighian tubules from third instar larvae of the sun beetle, *Pachnoda marginata*. In all images anterior is to the left. **(A,B)** Panels together forming an overview of the gut. Ventral view, showing foregut (fg), three rows of caeca (black arrows) in the midgut (mg), convoluted MpTs on the midgut surface (magenta arrowhead), the fermentation chamber (fc), and the seam between the two infoldings of the fermentation chamber (blue arrowhead). **(C)** Dorsal view of midgut with three rows of caeca indicated (white arrows) along with an additional pair of caeca between the anterior and central rows (blue arrow). **(D)** Ventral view. Upon pulling apart the folded sides of the fermentation chamber, and lifting back the rectum (r), four sinuous MpTs are revealed (arrowheads). **(E)** Unfolding the hindgut reveals the narrow ileum which had been obscured by the fermentation chamber. **(F)** Ventro-lateral view. Showing the insertion point of one of the laterally inserted pair of MpTs (blue arrowhead), one of the pair of MpTs which insert together on the ventral side of the gut (white arrowhead) and the midgut-hindgut boundary (white arrow).

**Figure 12.**
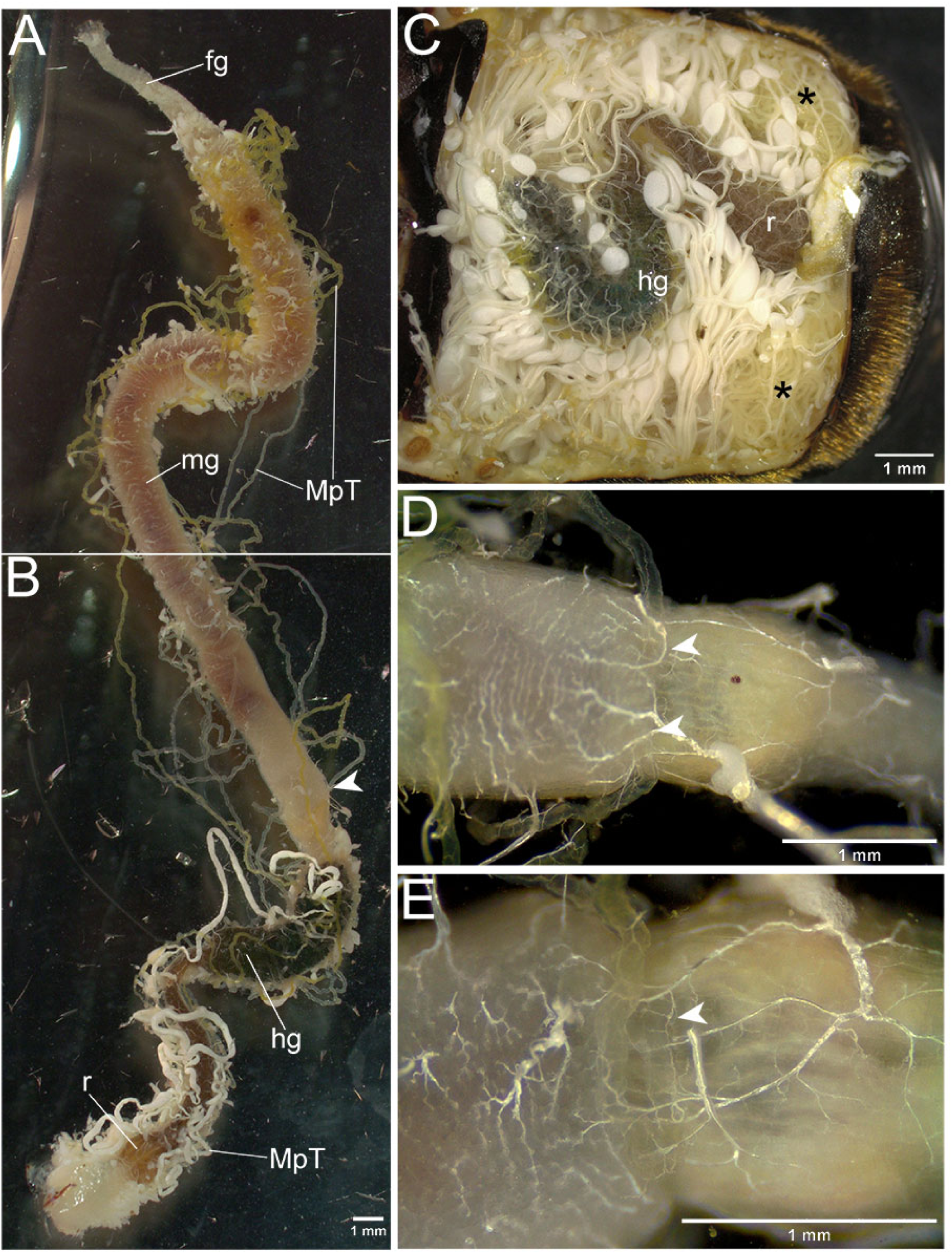
Gut and Malpighian tubules from adult sun beetles, *Pachnoda marginata*. **(A,B)** Dorsal view with anterior at the top, showing foregut (fg), midgut (mg), hindgut (hg), rectum (r) and midgut-hindgut boundary (arrow). **(C)** Showing position of gut in the body cavity, labelled as in B, and part of the male reproductive organs indicated (*). **(D)** A pair of MpTs insert into the gut in close proximity (arrowheads). **(E)** One of a pair of MpTs which insert laterally into the gut. Anterior is to the left in C-E.

## 4. Discussion

### 4.1 The independent evolution of a rectal complex in Scarabaeoidea

In arthropods, digestive and renal organs form part of a single integrated system, which underpins acquisition of nutrients, water and ionic homeostasis, and the removal of nitrogenous waste. They play a central role in survival, and have been modified at a molecular as well as morphological level during evolution, enabling shifts in diet and habitat. They have therefore played a critical role in the remarkable success of many arthropod groups.

The cryptonephridial or rectal complexes are one example of the evolutionary reorganisation of the arthropod digestive/renal system, and are considered to act as powerful recycling systems of water or solutes (Beaven et al., 2023; Kolosov and O’Donnell, 2019). The importance of rectal complexes is indicated by the multiple times they have evolved, and their prevalence, existing in several hundred thousand species, mainly within Lepidoptera and Coleoptera. The emerging picture from Scarabaeoidea, from assessing the phylogenetic distribution of rectal complexes observed in this and previous studies, is that a rectal complex is likely to have evolved independently within this group. Rectal complexes are restricted to part of the Scarabaeoidea tree, and are absent in more basal groups of Scarabaeoidea, and in the related group Staphylinoidea. Furthermore the structural differences we see in the rectal complexes of Scarabaeoidea, compared to those of Cucujiformia and Bostrichoidea, are also consistent with an independent evolutionary origin. Within Scarabaeoidea, rectal complexes are seemingly more variable between species, and plastic through insect life stages, than the rectal or cryptonephridial complexes found in Cucujiformia and Bostrichoidea. Rectal complexes in the Scarabaeoidea superfamily are therefore fertile ground for understanding the evolution and functions of these organ systems.

### 4.2 Structure of Scarabaeoidea rectal complexes and remodelling in pupal stage

Dramatic remodelling of the gut during pupal metamorphosis has been documented in *O. rhinoceros* (subfamily Dynastinae), as well as *Thaumastopeus shangaicus* and *Protaetia* species (subfamily Cetoniinae), which are all members of the Scarabaeidae family. It has been proposed that this is an adaptation to their radically different larval and adult diets (Chiang and Shelomi, 2023). Our observations show that there are also dramatic reorganisations of the MpTs and their relationship to the gut in Scarabaeoidea. In the case of *M. melolontha*, the rectal complex is likely to form during pupal metamorphosis. This is very different from the situation in tenebrionid beetles, where the complex is formed during embryonic development and appears to maintain a largely unchanged morphology between larval and adult stages (King and Denholm, 2014; Ramsay, 1964), however developmental remodelling of rectal complexes does occur in Lepidoptera, where they have only been observed in the larval stage (Barbehenn and Kristensen, 2003). The rectal complexes that we found in beetles from Cetoniinae are present in the larvae, and it remains to be determined whether they are established during embryonic or post-embryonic development.

We also saw evidence of major remodelling of the MpT regions not associated with the rectum, particularly in the occurrence of the branched form of MpT in adult *M. melolontha*. The role of this curious morphology is unclear. The branching would dramatically increase the surface area of the tubule available for molecular transport. These regions tend to associate with the midgut and hindgut surface, and branching could be to maximise coverage of the gut surface. Branches from the main MpT axis could also create a distinct microenvironment, potentially serving a number of purposes. The MpT branches are reminiscent of the diverticula or regenerative crypts seen in the midguts of some beetles such as adult *Tribolium*. These harbour stem cells which replenish the epithelium (Ameen and Rahman, 1973; Parthasarathy and Palli, 2008; Sinha, 1958; Snodgrass, 1994). Similar structures are seen in diverse arthropod groups including arachnids (Lipovšek et al., 2014; Lipovšek et al., 2022; Lipovšek et al., 2018). Bugs of the superfamilies Coreoidea and Lygaeoidea have structures termed caeca or crypts at the posterior end of their midguts, which appear to provide microenvironments to harbour symbiotic bacteria (Glasgow, 1914; Itoh et al., 2019; Takeshita and Kikuchi, 2020; Takeshita et al., 2015).

Differences can be seen between the structure of the rectal complexes of larval Cetoniinae and adult *M. melolontha* (Melolonthini). In larval Cetoniinae, we observed that sinuous MpTs coat the surface of the rectum, and are themselves enveloped within a fold in the hindgut fermentation chamber. It could therefore be considered that they from a unique structure: a hindgut fermentation chamber complex. The posterior rectum being enclosed by infoldings of the fermentation chamber has also been noted in larval *Protaetia* species and *Thaumastopeus shangaicus* (Cetoniinae) (Chiang and Shelomi, 2023), and in the case of *Protaetia cuprea*, sinuous MpTs surrounding the rectum have been described (Werner, 1926), consistent with our findings.

In *M. melolontha* adults there are also sinuous MpTs surrounding the rectum, but in this case we observe them to be surrounded by a membrane tissue, as has been previously reported (Lison, 1938; Saini, 1964). For the perinephric membrane, which has been characterised in most detail in *T. molitor* (Grimstone et al., 1968), the tissue entirely coats the CNC, except for small windows overlying specialised MpT cells (the leptophragmata) where it is reduced to blisters of extracellular material. In contrast, our observations suggest that the rectal complex membrane of *M. melolontha* is far from a continuous sheet, and may serve a structural role, rather than insulating the complex from the haemolymph as in tenebrionids, where it is considered to prevent the aberrant movement of water and ions. The differences we see in the rectal complexes between Melolonthini adults and Cetoniinae larvae make it conceivable that rectal complexes have evolved more than once within Scarabaeoidea, and future analysis of other groups should help clarify rectal complex evolutionary distribution and variability.

### 4.3 Putative functions of Scarabaeoidea rectal complexes: recycling water

In Tenebrionoidea the CNC appears to primarily function in water conservation. There is currently little evidence for whether rectal complexes fulfil a similar role in Scarabaeoidea. Within Lucanidae, which lack rectal complexes, there are indications of a requirement for very specific microhabitat conditions, including relatively high humidity (Scaccini, 2016; Scaccini, 2022). Further investigation will be required to determine the relative ability of groups with and without rectal complexes to retain water, and to tolerate periods of lower humidity. The rectum is an important site of water reabsorption, even in insect groups which lack rectal complexes (Phillips, 1964; Phillips et al., 1987; Ramsay, 1950; Ramsay, 1955; Wigglesworth, 1932; Wigglesworth, 1974). In larvae of some Scarabaeoidea species, the rectal contents can be expelled as a startle reflex, and such contents can be very wet and unformed. This is the case in species of Cetoniinae (*Cremastocheilus armatus* and *P. marginata*) as well as Lucanidae (*Platycerus* species, *D. parallelipipedus* and *L. cervus*) (Puker et al., 2015; our observations). These findings support the notion that drying of the gut contents would normally occur in the recta of larvae both with and without rectal complexes, and it remains to be clarified whether a rectal complex enhances this ability.

### 4.4 Putative functions of Scarabaeoidea rectal complexes: accessing nitrogen

A second possible function of rectal complexes in Scarabaeoidea relates to the availability of dietary nitrogen. For some termite species which have a diet very low in protein, there is evidence they utilise hindgut bacteria for recycling of nitrogen from excreted uric acid, and fixation of N_2_ (Benemann, 1973; Breznak et al., 1973; Desai and Brune, 2012; Ohkuma et al., 1999; Ohkuma et al., 1996; Potrikus and Breznak, 1981; Yamada et al., 2007).

Scarabaeoid beetles face similar challenges in accessing nitrogen, and nitrogen fixation has also been reported including in larval *Dorcus rectus* (Lucanidae) (Kuranouchi et al., 2006), and adult *Odontotaenius disjunctus* (Passalidae) (Ceja-Navarro et al., 2014). The contribution of nitrogen fixation to the nitrogen budget of *D. rectus* has been disputed (Tanahashi et al., 2018; our unpublished results), however recent findings from *Ceruchus piceus* (also from Lucanidae) support high levels of nitrogen fixation, which appears to be sensitive to experimental conditions (Mifsud et al., 2023). Interestingly the gut microbiome of *O. rhinoceros*, which is a beetle from Scarabaeoidea that feeds on rotting wood, is reported to be similar to that of wood feeding termites (Shelomi et al., 2019).

As insect MpTs act as transport sites for nitrogenous molecules including ammonia (Browne and O’Donnell, 2013), it is conceivable that their association with the rectum, in some groups of Scarabaeoidea, functions to transport ammonia from the gut contents into the body. Such a role could also point to the reason a rectal complex has been incorporated into the larval hindgut fermentation chamber, as this is the major site harbouring symbiotic bacteria (i.e. the presumptive site of nitrogen fixation or conversion of nitrogenous waste). This would be similar to the reported absorption of ammonia from the midgut contents demonstrated in Lepidoptera, which is considered to be an adaptation allowing rapid growth on a low protein diet (Blaesse et al., 2010; Hirayama et al., 1996; Weihrauch, 2006).

Some of the more basal groups of Scarabaeoidea such as Lucanidae larvae, lack rectal complexes. They are likely to have evolved distinct solutions to their dietary challenges, such as obtaining nitrogen and recycling water or solutes, as although absorption would likely occur in their rectum, this would be expected to be less efficient than in species with rectal complexes. For example they may use their environment as a kind of extended gut, enhancing their access to nitrogen and/or their ability to recycle water or ions. Supporting this notion, larvae of *D. rectus* and *L. cervus* eat decaying wood, and may re-ingest their own faeces (Hendriks and Fremlin, 2017; Tanahashi et al., 2018), whilst Cetoniinae larvae do not re-ingest their faeces (Tanahashi and Fremlin, 2013). There is also evidence that the fungi growing on dead wood could be an important dietary component for larvae from Lucanidae (Mishima and Araya, 2016; Tanahashi and Kubota, 2013; Tanahashi et al., 2009), with some species associating with wood of a specific decay type (Fukasawa, 2021; Scaccini, 2022). The associated fungi would be more easily digested, and likely provide more usable nitrogen, than wood alone (Tanahashi et al., 2009).

### 4.5 Putative functions of Scarabaeoidea rectal complexes: recycling bases

A third possibility is that the rectal complexes of Scarabaeoidea function in recycling bases from the gut contents. There is considerable variation in the reported midgut content pH in Scarabaeoidea species (Table 1; Huang et al., 2010), with some being very alkaline, for example larvae of *O. nasicornis* (Dynastinae) at pH 12.2 (Table 1). In this case the alkalinity has been proposed to function in precellulolysis, or chemically modifying cellulose to render it accessible to subsequent enzymatic breakdown in the hindgut (Bayon, 1980). In larval *Pachnoda ephippiata* (Cetoniinae) which have alkaline midgut contents (pH 10.7, Table 1), there is evidence that macromolecules, particularly humic acid bound peptides and polysaccharides, are rendered soluble in the midgut. This enables their depolymerisation by enzymes (Li and Brune, 2005a; Li and Brune, 2005b; Li and Brune, 2007). A highly alkaline midgut environment is also thought to enable the enzymatic activity of certain proteinases which can release amino acids from soil humic acid, as suggested by studies in this species (Zhang and Brune, 2004). High midgut pH may also be an adaptation to dietary tannins, although the possible effects of tannins on insects, such as inhibiting feeding, toxic effects or inhibiting digestion, has been a long contested issue (Ayres et al., 1997; Barbehenn and Constabel, 2011; Barbehenn et al., 2009a; Barbehenn et al., 2009b; Berenbaum, 1980; Bernays, 1978; Bernays et al., 1980; Bernays and Chamberlain, 1980; Bernays et al., 1981; Feeny, 1970; Feeny, 1968; Forkner et al., 2004; Karowe, 1989; Klocke and Chan, 1982; Martin et al., 1985; Rossiter et al., 1988; Tan et al., 2022).

**Table 1.** Diet, midgut pH and presence of rectal complex reported for members of the superfamily Scarabaeoidea. For midgut pH column, value is represented by colour, from green (mildly alkaline) to purple (highly alkaline). For rectal complex presence column, green indicates presence and magenta, absence. Ordered as in the phylogenetic tree in Figure 1.

| species | stage | diet | reported midgut pH | rectal complex presence |
| --- | --- | --- | --- | --- |
| *Dorcus rectus* (Lucanidae) | larva | rotting wood and associated fungi (Tanahashi and Kubota, 2013) | 10.2–10.4 (Tanahashi and Hawes, 2016) or 9.9 (Mishima and Araya, 2016) | unlikely |
| *Odontotaenius disjunctus* (Passalidae) | adult | rotting wood | 8.38 (Ceja-Navarro et al., 2014) | unlikely |
| *Sericesthis geminata* (Scitalini, part of Liparetrini in Fig. 1) | larva | roots | 9.0 (Soo Hoo and Dudzinski, 1967) | ? |
| *Costelytra zealandica* (Liparetrini) | larva | roots | 10.8 (Biggs and McGregor, 1996) | ? |
| *Melolontha melolontha* (Melolonthini) | larva | roots | 8.2 (Egert et al., 2005) | no (this study) |
| *Pachnoda marginata* (Cetoniinae) | larva | rotting leaves | 11.7 (Cazemier et al., 1997) | yes (this study) |
| *Pachnoda ephippiata* (Cetoniinae) | larva | rotting leaves | 10.7 (Lemke et al., 2003) | likely |
| *Oryctes nasicornis* (Dynastinae) | larva | rotting wood | 12.2 (Bayon, 1980) | yes (Bayon, 1981; Gérard, 1942; Sirodot, 1858) |
| *Trypoxylus dichotomus* (Dynastinae) | larva | rotting wood and leaves | 10.7 (Wada et al., 2014) | ? |

In Lepidoptera, high midgut pH is generated by transport of potassium and bicarbonate ions into the lumen (Dow, 1992), and their CNCs, which appear to have independently evolved, are thought to function in reabsorption of these ions downstream in the ileum and rectum. This would allow the larvae to maintain their acid-base balance (Kolosov and O’Donnell, 2019; Moffett, 1994; Ramsay, 1976).

A function of recycling ions also points to a possible explanation for why a rectal complex is present in Scarabaeoidea in some species and life stages, and absent in others. From the current reports existing for Scarabaeoidea, the most alkaline midguts are from larvae with a rectal complex, whilst larval *M. melolontha* (which we find to lack a rectal complex) has a less alkaline midgut (Table 1). This may relate to dietary properties, such as how resistant their food is to digestion, or how rich it is in tannin. The rotting leaves and wood eaten by many Cetoniinae larvae (Lemke et al., 2003; Ritcher, 1958), and the tree leaves eaten by adult *M. melolontha* (Ritcher, 1958), may be high in tannin, which could underly the presence of rectal complexes in these beetle species/life stages. On the other hand, the fruit or pollen and nectar eaten by Cetoniinae adults (Karolyi et al., 2009; Šípek et al., 2016), would contain much less tannin, which could have allowed evolutionary loss of the rectal complex in adulthood. *M. melolontha* larvae, which lack a rectal complex, feed on roots (Ritcher, 1958) which may have a lower tannin content than the leaves eaten by adults. There are some indications that tannin levels tend to be higher in leaves than roots (Dettlaff et al., 2018; Zhang et al., 2020).

Adoption of phytophagy (eating leaves) by insects is considered an evolutionary hurdle (Mitter et al., 1988; Southwood, 1972; Strong et al., 1984), that once surmounted has enabled major species radiations. Although rectal complexes or CNCs have been found in insect species with very diverse diets, it is noteworthy that many of the major evolutionary radiations of phytophagous insects appear to have occurred in groups with rectal complexes/CNCs. These include Phytophaga in the Cucujiformia infraorder of beetles (Crowson, 1981; Dufour, 1824; Farrell, 1998; Saini, 1964), and within Lepidoptera (Barbehenn and Kristensen, 2003; Powell et al., 1998). There has also been a significant radiation of phytophagous beetles in the Pleurosticti lineage within Scarabaeoidea (Ahrens et al., 2014; Fig. 1). More extensive sampling of digestive/renal system anatomies could inform whether rectal complexes are likely to have contributed to the adoption of phytophagy within Scarabaeoidea.

Lucanidae larvae may also be adapted to digest highly resistant materials such as rotting wood, and also have a fairly high midgut pH of ∼10.3 (Table 1). They are likely to have evolved alternative strategies, such as re-ingestion of faeces, in order to efficiently recover bases from their gut contents.

### 4.6 Ecological implications of digestive/renal systems of Scarabaeoidea

Scarabaeoidea is a very large beetle superfamily, comprising ∼35,000 species (Hunt et al., 2007). These beetles play critical roles in many ecosystems, for example in the breakdown of resistant plant materials, such as dung and wood, rendering their components accessible in the soil (Stanbrook et al., 2021; Stokland, 2012; Ulyshen, 2018). These functions must rely on adaptations of their digestive/renal systems. The ability of Scarabaeoidea species to digest highly resistant material such as lignocellulose is central to this, and relies on the symbiotic bacteria and yeasts within their guts (Ebert et al., 2021; Mishima and Araya, 2016; Shelomi et al., 2019; Tanahashi et al., 2017; Vargas-Asensio et al., 2014). For this reason *Protaetia brevitarsis*, a member of Cetoniinae farmed as a food source in East Asia, is being investigated for its ability to convert organic waste into fertiliser (Li et al., 2019; Wang et al., 2022). There is evidence from *Pachnoda ephippiata* (Cetoniinae) that their fresh faeces are rich in ammonia, which is a usable nitrogen source for plants (Li and Brune, 2007). Whilst microbial activity and the availability of usable nutrients is high in fresh faecal pellets of several Cetoniinae species, this may be relatively short lived; the older pellets are highly resistant to further degradation, and may then serve other ecosystem services such as sequestration of carbon (Jönsson et al., 2004; Kulikova and Perminova, 2021; Li and Brune, 2007; Micó et al., 2011; Sánchez et al., 2017). The faeces of Cetoniinae larvae can remain undegraded for many years (Jönsson et al., 2004; Martin, 1997; Pawlowski, 1961; Ranius and Nilsson, 1997; our observations). There is also evidence that Scarabaeoidea larvae can act as ecosystem engineers, for example the presence of *Cetonia aurataeformis* appears to benefit the larvae of hover fly species, also living on rotting wood (Sánchez-Galván et al., 2014). Shedding light on the digestive/renal systems found in Scarabaeoidea is important for understanding the ecological roles played by its members.

### 4.7 Conclusion and future directions

This study is an initial exploration of the presence and morphology of rectal complexes in beetles of the Scarabaeoidea superfamily. Further sampling of the basal groups of Scarabaeoidea, along with the related superfamilies Staphylinoidea and Hydrophiloidea (Fig.1 and S1) could provide further evidence of whether this complex evolved independently in Scarabaeoidea. More extensive studies comparing larvae and adults in species across the evolutionary tree of Scarabaeoidea, could yield stronger evidence of when a rectal complex evolved within Scarabaeoidea, and differences in morphology between groups could also illuminate the course of its evolution. By relating to ecological aspects of each species and life stage, such as diet and environmental conditions, and physiological data such as gut pH, this could shed further light on the evolutionary significance of rectal complexes in Scarabaeoidea. It is possible that the evolutionary acquisition of rectal complexes, as well as their secondary losses, have been governed by a combination of the functions outlined above. Ultimately unpicking their precise functional roles will require targeted molecular approaches, in combination with physiological studies.

## Acknowledgements

We are extremely grateful to David Beaven for sending a *M. melolontha* adult, and Jordan Johnson and Rebecca Kilner for sending *N. vespilloides* larvae. We also thank Anisha Kubasik-Thayil and Adrian Garcia Burgos from the IMPACT imaging facility, University of Edinburgh. This work was supported by The Leverhulme Trust Grant (RPG-2019-167), the BBSRC (BB/X014703/1), a Moray Endowment Fund Award (University of Edinburgh) and the Deanery of Biomedical Sciences (University of Edinburgh). For the purpose of open access, the authors have applied a CC BY public copyright licence to any Author Accepted Manuscript version arising from this submission.

---

**Figure S1.**
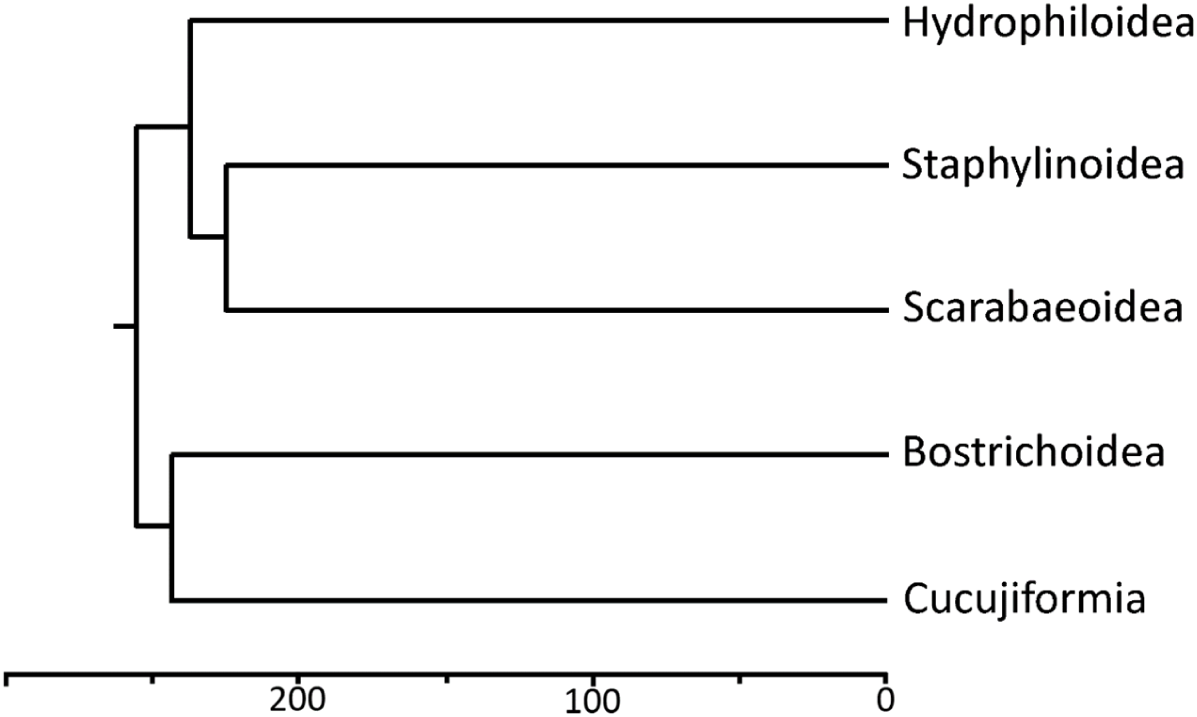
Phylogenetic tree of Polyphaga suborder of beetles. CNCs occur in Bostrichoidea and Cucujiformia. The tree is modified from McKenna et al., 2019, with estimated evolutionary time shown (millions of years). In the tribe Melolonthini, apart from the rectal complex seen in *M. melolontha* adults, they were also reported to be present in *Phyllophaga anxia* larvae and adults (Berberet and Helms, 1972), *Amphimallon majale* (Menees, 1958), and *Polyphylla decemitheata* larvae (Areekul, 1957), and *Phyllophaga gracilis* larvae (Areekul, 1957) but not adults (Fletcher, 1930). A rectal complex was not reported in larvae or adults of the tribe Rhizotrogini (Schäfer, 1954), or the subfamily Rutelinae (Nagae et al., 2013; Swingle, 1930), or adults of Diplotaxini (Jones, 1940). A rectal complex has been reported in larvae in the family Glaphyridae (Areekul, 1957). It was also reportedly present in larvae, but not adults, of the subfamily Cetoniinae (Dufour, 1824; Dufour, 1842; Werner, 1926). Within the subfamily Dynastinae, a rectal complex was observed in *Oryctes nasicornis* larvae (Bayon, 1981; Gérard, 1942; Sirodot, 1858), but not *Heteronychus aratar* larvae or adults (Sheehan et al., 1982). A putative evolutionary distribution of rectal complexes is shown in Figure 1. Note that a more recent phylogeny based on transcriptomic data gave somewhat different relationships, for example placing Glaphyridae in a more basal position (Dietz et al., 2023). The current picture is complex, and may be explained by multiple gains and/or losses of rectal complexes in Scarabaeoidea, although the limited sampling and variable quality of previous reporting adds to the uncertainty. A comprehensive understanding of the evolutionary distribution of rectal complexes, and their precise structure and functional role(s), remain unknown in Scarabaeoidea.

**Figure S2.**
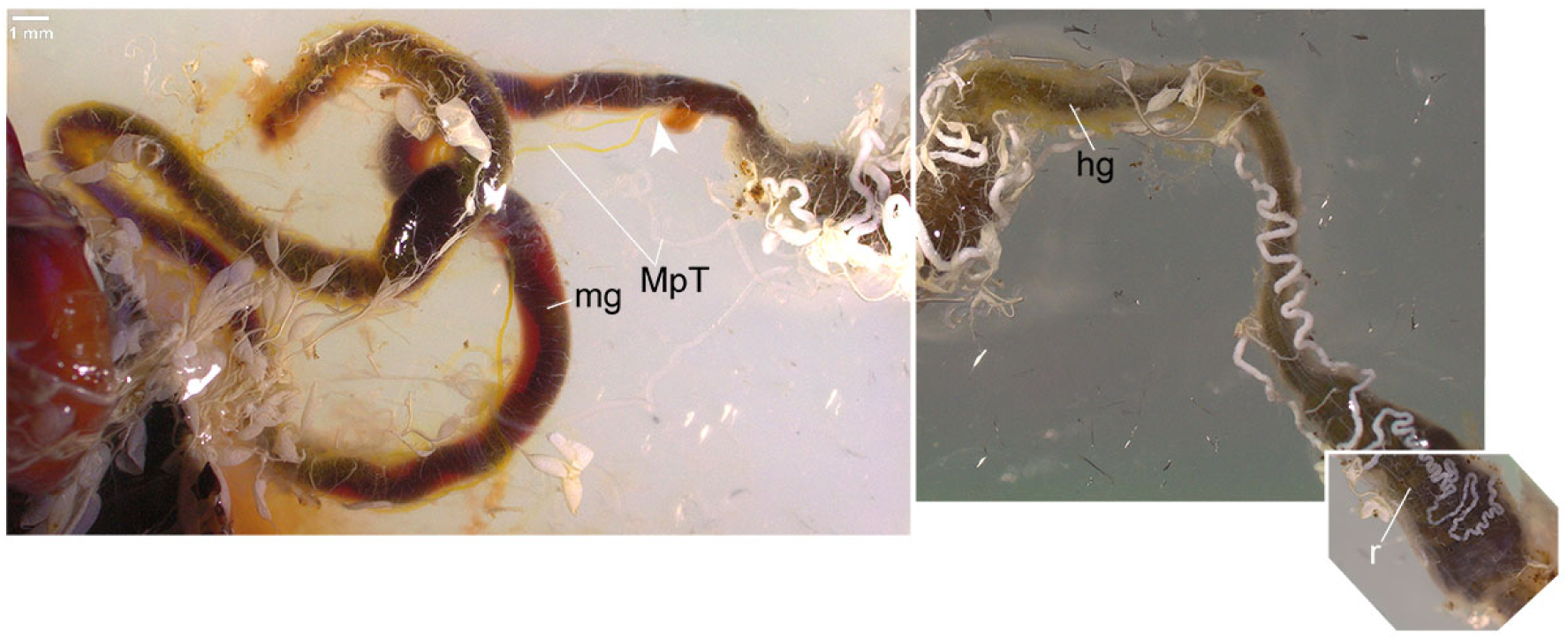
Gut and MpTs of an adult male cockchafer, *Melolontha melolontha*. Montage of images in which the midgut (mg) and hindgut (hg) are indicated, along with the boundary between the two (arrowhead), as well as the rectum (r). Note that the most anterior end of gut is missing, from dissection.

## Notes

### Competing Interest Statement

The authors have declared no competing interest.

## References

Ahrens, D., Schwarzer, J. and Vogler, A. P. (2014). The evolution of scarab beetles tracks the sequential rise of angiosperms and mammals. *Proceedings*. Biological sciences 281.

Ameen, M. and Rahman, M. F. (1973). Larval and adult digestive tracts of *Tribolium castaneum* (Herbst) (Coleoptera: Tenebrionidae). International Journal of Insect Morphology and Embryology 2, 137–152.

Arab, A. and Caetano, F. H. (2002). Segmental specializations in the Malpighian tubules of the fire ant *Solenopsis saevissima* Forel 1904 (Myrmicinae): an electron microscopical study. Arthropod Structure & Development 30, 281–292.

Areekul, S. (1957). The Comparative Internal Larval Anatomy of Several Genera of Scarabaeidae (Coleoptera). Annals of the Entomological Society of America 50, 562–577.

Ayres, M. P., Clausen, T. P., MacLean, S. F., Redman, A. M. and & Reichardt, P. B. (1997). Diversity of Structure and Antiherbivore Activity in Condensed Tannins. Ecology 78, 1696–1712.

Barbehenn, R. V. and Constabel, C. P. (2011). Tannins in plant-herbivore interactions. Phytochemistry 72, 1551–1565.

Barbehenn, R. V., Jaros, A., Lee, G., Mozola, C., Weir, Q. and Salminen, J. P. (2009a). Hydrolyzable tannins as “quantitative defenses”: limited impact against Lymantria dispar caterpillars on hybrid poplar. Journal of insect physiology 55, 297–304.

Barbehenn, R. V., Jaros, A., Lee, G., Mozola, C., Weir, Q. and Salminen, J. P. (2009b). Tree resistance to *Lymantria dispar* caterpillars: importance and limitations of foliar tannin composition. Oecologia 159, 777–788.

Barbehenn, R. V. and Kristensen, N. P. (2003). 6. Digestive and excretory systems. In Teilband/Part 36 Vol 2: Morphology, Physiology, and Development (ed. W. Kükenthal), pp. 165–188. Berlin: De Gruyter.

Bayon, C. (1980). Volatile fatty acids and methane production in relation to anaerobic carbohydrate fermentation in Oryctes nasicornis larvae (Coleoptera: Scarabaeidae). Journal of Insect Physiology 26, 819–828.

Bayon, C. (1981). Ultrastructure de l’epithelium intestinal et flore parietale chez la larve xylophage d’*Oryctes nasicornis* L. (Coleoptera : scrabaeidae). International Journal of Insect Morphology and Embryology 10, 359–371.

Beaven, R., Halberg, K. V. and Denholm, B. (2023). The insect cryptonephridial complex. Current biology 33, R748–R749.

Benemann, J. R. (1973). Nitrogen fixation in termites. Science 181, 164–165.

Berberet, R. C. and Helms, T. J. (1972). Comparative Anatomy and Histology of Selected Systems in Larval and Adult *Phyllophaga anxia* (Coleoptera: Scarabaeidae). Annals of the Entomological Society of America 65, 1026–1053.

Berenbaum, M. (1980). Adaptive Significance of Midgut pH in Larval Lepidoptera. The American Naturalist 115, 138–146.

Bernays, E. A. (1978). Tannins: An Alternative Viewpoint. Entomologia experimentalis et applicata 24, 244–253.

Bernays, E. A., Chamberlain, D. and McCarthy, P. (1980). The differential effects of ingested tannic acid on different species of Acridoidea. Entomologia Experimentalis et Applicata 28, 158–166.

Bernays, E. A. and Chamberlain, D. J. (1980). A study of tolerance of ingested tannin in *Schistocerca gregaria*. Journal of Insect Physiology 26, 415–420.

Bernays, E. A., Chamberlain, D. J. and Leather, E. M. (1981). Tolerance of acridids to ingested condensed tannin. Journal of chemical ecology 7, 247–256.

Beutel, R. G. and Haas, F. (2000). Phylogenetic Relationships of the Suborders of Coleoptera (Insecta). Cladistics : the international journal of the Willi Hennig Society 16, 103–141.

Biggs, D. R. and McGregor, P. G. (1996). Gut pH and amylase and protease activity in larvae of the New Zealand grass grub (*Costelytra zealandica*; Coleoptera: Scarabaeidae) as a basis for selecting inhibitors. Insect Biochemistry and Molecular Biology 26, 69–75.

Blaesse, A. K., Broehan, G., Meyer, H., Merzendorfer, H. and Weihrauch, D. (2010). Ammonia uptake in *Manduca sexta* midgut is mediated by an amiloride sensitive cation/proton exchanger: Transport studies and mRNA expression analysis of NHE7, 9, NHE8, and V-ATPase (subunit D). Comparative biochemistry and physiology. Part A, Molecular & integrative physiology 157, 364–376.

Breznak, J. A., Brill, W. J., Mertins, J. W. and Coppel, H. C. (1973). Nitrogen fixation in termites. Nature 244, 577–580.

Browne, A. and O’Donnell, M. J. (2013). Ammonium secretion by Malpighian tubules of *Drosophila melanogaster*: application of a novel ammonium-selective microelectrode. The Journal of experimental biology 216, 3818–3827.

Cazemier, A. E., Op den Camp, H. J., Hackstein, J. H. P. and Vogels, G. D. (1997). Fibre digestion in arthropods. Comparative Biochemistry and Physiology 118A, 101–109.

Ceja-Navarro, J. A., Nguyen, N. H., Karaoz, U., Gross, S. R., Herman, D. J., Andersen, G. L., Bruns, T. D., Pett-Ridge, J., Blackwell, M. and Brodie, E. L. (2014). Compartmentalized microbial composition, oxygen gradients and nitrogen fixation in the gut of *Odontotaenius disjunctus*. The ISME journal 8, 6–18.

Chapman, R. F. (1998). Alimentary canal, digestion and absorption. In The Insects: Structure and Function (ed. R. F. Chapman), pp. 38–68. Cambridge: Cambridge University Press.

Chiang, M. R. and Shelomi, M. (2023). Anatomical changes of the beetle digestive tract during metamorphosis correspond to dietary changes. Journal of morphology 284.

Coutchié, P. A. and Machin, J. (1984). Allometry of water vapor absorption in two species of tenebrionid beetle larvae. American Journal of Physiology 247, R230–236.

Crowson, R. A. (1981). Chapter 18 - Herbivorous Beetles. In The Biology of the Coleoptera, pp. 584–618: Academic Press.

Dallai, R., Del Bene, G. and Marchini, D. (1991). The ultrastructure of Malpighian tubules and hindgut of *Frankliniella occidentalis* (Pergande) (Thysanoptera : Thripidae). International Journal of Insect Morphology and Embryology 20, 223–233.

Denholm, B. (2013). Shaping up for action: the path to physiological maturation in the renal tubules of *Drosophila*. Organogenesis 9, 40–54.

Desai, M. S. and Brune, A. (2012). Bacteroidales ectosymbionts of gut flagellates shape the nitrogen-fixing community in dry-wood termites. The ISME journal 6, 1302–1313.

Dettlaff, M. A., Marshall, V., Erbilgin, N. and Cahill, J. F. (2018). Root condensed tannins vary over time, but are unrelated to leaf tannins. AoB PLANTS 10.

Dietz, L., Seidel, M., Eberle, J., Misof, B., Pacheco, T. L., Podsiadlowski, L., Ranasinghe, S., Gunter, N. L., Niehuis, O., Mayer, C., et al. (2023). A transcriptome-based phylogeny of Scarabaeoidea confirms the sister group relationship of dung beetles and phytophagous pleurostict scarabs (Coleoptera). Systematic Entomology 48, 672 – 686.

Dow, J. A. (1992). pH Gradients in Lepidopteran Midgut. The Journal of experimental biology 172, 355–375.

Dow, J. A., Maddrell, S. H., Gortz, A., Skaer, N. J., Brogan, S. and Kaiser, K. (1994). The Malpighian tubules of *Drosophila melanogaster*: a novel phenotype for studies of fluid secretion and its control. J Exp Biol 197, 421–428.

Dufour, M. L. (1824). Recherches anatomiques sur les Carabiques et sur plusieurs autres insectes Coléoptères. Annales des Sciences Naturelles Atlas des Tomes 1,2,3, 27–42.

Dufour, M. L. (1842). Histoire comparative des métamorphoses et de l’anatomie des *Cetonia aurata* et *Dorcus parallelipipedus*. Annales Des Sciences Naturelles 18, 162–181.

Dunbar, B. S. and Winston, P. W. (1975). The site of active uptake of atmospheric water in larvae of *Tenebrio molitor*. Journal of Insect Physiology 21, 495–500.

Ebert, K. M., Arnold, W. G., Ebert, P. R. and Merritt, D. J. (2021). Hindgut microbiota reflects different digestive strategies in dung beetles (Coleoptera: Scarabaeidae: Scarabaeinae). Applied and environmental microbiology 87, e02100–02120.

Edwards, E. E. (1930). On the Morphology of the Larva of *Dorcus parallelopipedus*, L. (Coleoptera). Zoological journal of the Linnean Society 37, 93–108.

Egert, M., Stingl, U., Bruun, L. D., Pommerenke, B., Brune, A. and Friedrich, M. W. (2005). Structure and topology of microbial communities in the major gut compartments of *Melolontha melolontha* larvae (Coleoptera: Scarabaeidae). Applied and environmental microbiology 71, 4556–4566.

Farrell, B. D. (1998). “Inordinate Fondness” Explained: Why Are There So Many Beetles? Science 281, 555–559.

Feeny, P. (1970). Seasonal Changes in Oak Leaf Tannins and Nutrients as a Cause of Spring Feeding by Winter Moth Caterpillars. Ecology 51, 565–581.

Feeny, P. P. (1968). Effect of oak leaf tannins on larval growth of the winter moth *Operophtera brumata*. Journal of Insect Physiology 14, 805–817.

Fletcher, F. W. (1930). The Alimentary Canal of *Phyllophaga Gracilis* Burm. The Ohio Journal of Science 30, 109–119.

Forkner, R. E., Marquis, R. J. and Lill, J. T. (2004). Feeny revisited: condensed tannins as anti-herbivore defences in leaf-chewing herbivore communities of *Quercus*. Ecological Entomology 29, 174–187.

Fremlin, M. (2018). The Rose Chafer C*etonia aurata* L. (Coleoptera: Scarabaeidae:Cetoniinae) in Essex: distribution and some aspects of its ecology. Essex Naturalist 35, 167–178.

Fukasawa, Y. (2021). Ecological impacts of fungal wood decay types: A review of current knowledge and future research directions. Ecological Research 36, 910 – 931.

Glasgow, H. (1914). The Gastric Cæca and the Cæcal Bacteria of the Heteroptera. Biological Bulletin 26, 101–170.

Grimstone, A. V., Mullinger, A. M. and Ramsay, J. A. (1968). Further studies on the rectal complex of mealworm *Tenebrio molitor*, L. (Coleoptera, Tenebrioidae). Philosophical Transactions of the Royal Society of London. Series B, Biological Sciences 253, 343–382.

Gérard, P. (1942). Les tubes de Malpighi de la larve d’*Oryctes Nasicornis* L. Annales de la Société royale zoologique de Belgique 73, 122–133.

Hansen, L. L., Ramløv, H. and Westh, P. (2004). Metabolic activity and water vapour absorption in the mealworm *Tenebrio molitor* L. (Coleoptera, Tenebrionidae): real-time measurements by two-channel microcalorimetry. Journal of Experimental Biology 207, 545–552.

Hendriks, P. and Fremlin, M. (2017). How stag beetle Lucanus cervus larvae feed. http://maria.fremlin.de/stagbeetles/larva_feeding/index.html.

Henson, H. (1937). The Structure and Post-Embryonic Development of *Vanessa urticse* (Lepidoptera)–II. The Larval Malpighian Tubules. Proceedings of the Zoological Society of London 107, 161–174.

Hirayama, C., Konno, K. and Shinbo, H. (1996). Utilization of ammonia as a nitrogen source in the silkworm, *Bombyx mori*. Journal of Insect Physiology 42, 983–988.

Huang, S.-W., Zhang, H.-Y., Marshall, S. and Jackson, T. A. (2010). The scarab gut: A potential bioreactor for bio-fuel production. Insect Science 17, 175–183.

Hunt, T., Bergsten, J., Levkanicova, Z., Papadopoulou, A., John, O. S., Wild, R., Hammond, P. M., Ahrens, D., Balke, M., Caterino, M. S., et al. (2007). A comprehensive phylogeny of beetles reveals the evolutionary origins of a superradiation. Science (New York, N.Y.) 318, 1913–1916.

Irvine, H. B. (1969). Sodium and potassium secretion by isolated insect Malpighian tubules. The American journal of physiology 217, 1520–1527.

Ito, H. (1921). On the metamorphosis of the Malpighian tubes of *Bombyx Mori* L. Journal of Morphology 35, 195–211.

Itoh, H., Jang, S., Takeshita, K., Ohbayashi, T., Ohnishi, N., Meng, X., Mitani, Y. and Kikuchi, Y. (2019). Host-symbiont specificity determined by microbe-microbe competition in an insect gut. Proceedings of the National Academy of Sciences of the United States of America 116, 22673–22682.

Jones, C. R. (1940). The alimentary canal of *Diplotaxis liberta* Germ. (Scarabaeidae: Coleoptera). The Ohio Journal of Science 40, 94–103.

Judy, K. (1968). Studies on the metamorphosis of the alimentary canal of Hyalophora cecropia (L.). ProQuest Dissertations Publishing.

Jönsson, N., Méndez, M. and Ranius, T. (2004). Nutrient richness of wood mould in tree hollows with the Scarabaeid beetle *Osmoderma eremita*. Animal Biodiversity and Conservation 27, 79–82.

Karolyi, F., Gorb, S. N. and Krenn, H. W. (2009). Pollen grains adhere to the moist mouthparts in the flower visiting beetle C*etonia aurata* (Scarabaeidae, Coleoptera). Arthropod-Plant Interactions 3, 1–8.

Karowe, D. N. (1989). Differential effect of tannic acid on two tree-feeding Lepidoptera: implications for theories of plant anti-herbivore chemistry. Oecologia 80, 507–512.

King, B. and Denholm, B. (2014). Malpighian tubule development in the red flour beetle (Tribolium castaneum). Arthropod Structure & Development 43, 605–613.

Kirschner, L. B. (1967). Comparative Physiology: Invertebrate Excretory Organs. Annual Review of Physiology 29, 169–196.

Klocke, J. A. and Chan, B. G. (1982). Effects of cotton condensed tannin on feeding and digestion in the cotton pest, *Heliothis zea*. Journal of Insect Physiology 28, 911–915.

Koefoed, B. (1971). Ultrastructure of the cryptonephridial system in the meal worm *Tenebrio molitor*. Zeitschrift für Zellforschung und Mikroskopische Anatomie 116, 487–501.

Kolosov, D. and O’Donnell, M. J. (2019). Chapter Five - The Malpighian tubules and cryptonephric complex in lepidopteran larvae. Advances in Insect Physiology 56, 165–202.

Kucuk, R. A., Campbell, B. J., Lyon, N. J., Shelby, E. A. and Caterino, M. S. (2023). Gut bacteria of adult and larval *Cotinis nitida* Linnaeus (Coleoptera: Scarabaeidae) demonstrate community differences according to respective life stage and gut region. Frontiers in microbiology 14.

Kulikova, N. A. and Perminova, I. V. (2021). Interactions between Humic Substances and Microorganisms and Their Implications for Nature-like Bioremediation Technologies. Molecules 26.

Kuranouchi, T., Nakamura, T., Shimamura, S., Kojima, H., Goka, K., Okabe, K. and Mochizuki, A. (2006). Nitrogen fixation in the stag beetle, *Dorcus* (*Macrodorcus*) *rectus* (Motschulsky) (Col., Lucanidae). Journal of Applied Entomology 130, 471–472.

Lawrence, J. F. and Newton, A. F. (1982). Evolution and Classification of Beetles. Annual Review of Ecology and Systematics 13, 261–290.

Lemke, T., Stingl, U., Egert, M., Friedrich, M. W. and Brune, A. (2003). Physicochemical conditions and microbial activities in the highly alkaline gut of the humus-feeding larva of *Pachnoda ephippiata* (Coleoptera: Scarabaeidae). Applied and environmental microbiology 69, 6650–6658.

Li, X. and Brune, A. (2005a). Digestion of microbial biomass, structural polysaccharides, and protein by the humivorous larva of *Pachnoda ephippiata* (Coleoptera: Scarabaeidae). Soil Biology and Biochemistry 37, 107–116.

Li, X. and Brune, A. (2005b). Selective digestion of the peptide and polysaccharide components of synthetic humic acids by the humivorous larva of *Pachnoda ephippiata* (Coleoptera: Scarabaeidae). Soil Biology and Biochemistry 37, 1476–1483.

Li, X. and Brune, A. (2007). Transformation and mineralization of soil organic nitrogen by the humivorous larva of *Pachnoda ephippiata* (Coleoptera: Scarabaeidae). Plant and Soil 301, 233–244.

Li, Y., Fu, T., Geng, L., Shi, Y., Chu, H., Liu, F., Liu, C., Song, F., Zhang, J. and Shu, C. (2019). *Protaetia brevitarsis* larvae can efficiently convert herbaceous and ligneous plant residues to humic acids. Waste management (New York, N.Y.) 83, 79–82.

Lipovšek, S., Janžekovič, F. and Novak, T. (2014). Autophagic activity in the midgut gland of the overwintering harvestmen *Gyas annulatus* (Phalangiidae, Opiliones). Arthropod structure & development 43, 493–500.

Lipovšek, S., Novak, T., Dariš, B., Hofer, F., Leitinger, G. and Letofsky-Papst, I. (2022). Ultrastructure of spherites in the midgut diverticula and Malpighian tubules of the harvestman *Amilenus aurantiacus* during the winter diapause. Histochemistry and cell biology 157, 107–118.

Lipovšek, S., Novak, T., Janžekovič, F., Brdelak, N. and Leitinger, G. (2018). Changes in the midgut diverticula epithelial cells of the European cave spider, *Meta menardi*, under controlled winter starvation. Scientific Reports 8, 1–13.

Lison, L. (1937). Sur la structure de la region cryptosoleniee chez les Coleopteres *Tenebrio molitor* L. et *Dermestes lardarius* L. Bulletin de la Classe des Sciences, Académie Royale de Belgique/Mededeelingen van de Afdeeling Wetenschappen, Koninklije Belgische Academie 23, 317–327.

Lison, L. (1938). Contribution à l’étude morphologique et histophysiologique du système malpighien de M*elolontha melolontha* Linn. (Coleoptera). Annales de la Société royal zoologique de Belgique 69, 195–233.

Machin, J. (1975). Water balance in *Tenebrio molitor*, L. Larvae; the effect of atmospheric water absorption. Journal of comparative physiology 101, 121–132.

Machin, J. (1979). Compartmental Osmotic Pressures in the Rectal Complex of *Tenebrio* Larvae: Evidence for a Single Tubular Pumping Site. The Journal of Experimental Biology 82, 123–137.

Machin, J. and O’Donnell, M. J. (1991). Rectal complex ion activities and electrochemical gradients in larvae of the desert beetle, *Onymacris*: Comparisons with *Tenebrio*. Journal of Insect Physiology 37, 829–838.

Marshall, A. T. and Wright, A. (1972). Detection of diffusible ions in insect osmoregulatory systems by electron probe X-ray microanalysis using scanning electron microscopy and a cryoscopic technique. Micron (1969) 4, 31–45.

Martin, M. M., Rockholm, D. C. and Martin, J. S. (1985). Effects of surfactants, pH, and certain cations on precipitation of proteins by tannins. Journal of Chemical Ecology 11, 485–494.

Martin, O. (1997). Fredede insekter i Danmark. Del 2: Biller knyttet til skov. Entomologiske Meddelelser 61, 63 – 76.

Menees, J. H. (1958). The Anatomy and Histology of the Larval Alimentary Canal of the European Chafer, *Amphimallon Majalis* Razoumowsky (Scarabaeidae). Journal of the New York Entomological Society 66, 75–86.

Metalnikov, S. (1908). Recherches expérimentales sur les Chenilles de *Galleria mellonella*. Archives de zoologie expérimentale et générale 4, 489–588.

Micó, E., Juárez, M., Sánchez, A. and Galante, E. (2011). Action of the saproxylic scarab larva *Cetonia aurataeformis* (Coleoptera: Scarabaeoidea: Cetoniidae) on woody substrates. Journal of Natural History 45, 41–42.

Mifsud, I. E. J., Akana, P. R., Bytnerowicz, T. A., Davis, S. R. and Menge, D. N. L. (2023). Nitrogen fixation in the stag beetle, *Ceruchus piceus* (Coleoptera: Lucanidae): could insects contribute more to ecosystem nitrogen budgets than previously thought? Environmental entomology 52, 618–626.

Mishima, T. and Araya, K. (2016). Are the larvae of stag beetles xylophagous or mycophagous? - Analysis of polysaccharide digestive enzymatic systems of the larvae of *Dorcus rectus rectus*. Kogane 18, 94–100.

Mitter, C., Farrell, B. and Wiegmann, B. (1988). The Phylogenetic Study of Adaptive Zones: Has Phytophagy Promoted Insect Diversification? The American Naturalist 132, 107–128.

Moffett, D. F. (1994). Recycling of K⁺, Acid-Base Equivalents, and Fluid between Gut and Hemolymph in Lepidopteran Larvae. Physiological Zoology 67, 68–81.

Nagae, T., Miyake, S., Kosaki, S. and Azuma, M. (2013). Identification and characterisation of a functional aquaporin water channel (*Anomala cuprea* DRIP) in a coleopteran insect. The Journal of Experimental Biology 216, 2564–2572.

Naseem, M. T., Beaven, R., Koyama, T., Naz, S., Su, S. Y., Leader, D. P., Klaerke, D. A., Calloe, K., Denholm, B. and Halberg, K. V. (2023). NHA1 is a cation/proton antiporter essential for the water-conserving functions of the rectal complex in *Tribolium castaneum*. Proceedings of the National Academy of Sciences of the United States of America 120.

Noble-Nesbitt, J. (1970). Water uptake from subsaturated atmospheres: its site in insects. Nature 225, 753–754.

O’Donnell, M. J. and Machin, J. (1991). Ion Activities and Electrochemical Gradients in the Mealworm Rectal Complex. Journal of Experimental Biology 155, 375–402.

O’Donnell, M. J. and Ruiz-Sanchez, E. (2015). The rectal complex and Malpighian tubules of the cabbage looper (*Trichoplusia ni*): regional variations in Na+ and K+ transport and cation reabsorption by secondary cells. The Journal of Experimental Biology 218, 3206.

Ohkuma, M., Noda, S. and Kudo, T. (1999). Phylogenetic diversity of nitrogen fixation genes in the symbiotic microbial community in the gut of diverse termites. Applied and environmental microbiology 65, 4926–4934.

Ohkuma, M., Noda, S., Usami, R., Horikoshi, K. and Kudo, T. (1996). Diversity of Nitrogen Fixation Genes in the Symbiotic Intestinal Microflora of the Termite *Reticulitermes speratus*. Applied and environmental microbiology 62, 2747–2752.

Parthasarathy, R. and Palli, S. R. (2008). Proliferation and differentiation of intestinal stem cells during metamorphosis of the red flour beetle, *Tribolium castaneum*. Developmental dynamics : an official publication of the American Association of Anatomists 237, 893–908.

Pawlowski, J. (1961). Próchnojady blazkorozne w biocenozie leśnej Polski – Lamellicornes cariophages in forest biocenosis of Poland. Ekologia Polska Seria A 9, 355–437.

Phillips, J. E. (1964). Rectal Absorption in the Desert Locust, *Schistocerca Gregaria* Forskål I. Water. Journal of Experimental Biology 41, 15–38.

Phillips, J. E. (1970). Apparent Transport of Water by Insect Excretory Systems. American Zoologist 10, 413–436.

Phillips, J. E., Thomson, B., Hanrahan, J. and Chamberlin, M. (1987). Mechanisms and Control of Reabsorption in Insect Hindgut. Advances in Insect Physiology 19, 329–422.

Polilov, A. A. (2008). Anatomy of the smallest coleoptera, featherwing beetles of the tribe nanosellini (Coleoptera, Ptiliidae), and limits of insect miniaturization. Entomological Review 88, 26–33.

Potrikus, C. J. and Breznak, J. A. (1981). Gut bacteria recycle uric acid nitrogen in termites: A strategy for nutrient conservation. Proceedings of the National Academy of Sciences of the United States of America 78, 4601–4605.

Powell, J. A., Mitter, C. and Farrell, B. (1998). 20. Evolution of Larval Food Preferences in Lepidoptera. In Teilband/Part 35 Volume 1: Evolution, Systematics, and Biogeography (ed. W. Kükenthal), pp. 403–422. Berlin, Boston: De Gruyter.

Puker, A., Rosa, C. S., Orozco, J., Solar, R. R. C. and Feitosa, R. M. (2015). Insights on the association of American Cetoniinae beetles with ants. Entomological Science 18, 21–30.

Ramsay, J. A. (1950). Osmotic regulation in mosquito larvae. The Journal of Experimental Biology 27, 145.

Ramsay, J. A. (1955). The excretory system of the stick insect, *Dixippus morosus* (Orthoptera, Phasmidae). J. exp. Biol. 32, 183 – 199.

Ramsay, J. A. (1964). The rectal complex of the mealworm *Tenebrio molitor*, L. (Coleoptera, Tenebrionidae). Philosophical Transactions of the Royal Society of London. Series B, Biological Sciences 248, 279–314.

Ramsay, J. A. (1976). The rectal complex in the larvae of Lepidoptera. *Philosophical Transactions of the Royal Society of London*. B, Biological Sciences 274, 203–226.

Ranius, T. and Nilsson, S. G. (1997). Habitat of *Osmoderma eremita* Scop. (Coleoptera: Scarabaeidae), a beetle living in hollow trees. Journal of Insect Conservation 1, 193–204.

Regier, J. C., Zwick, A., Cummings, M. P., Kawahara, A. Y., Cho, S., Weller, S., Roe, A., Baixeras, J., Brown, J. W., Parr, C., et al. (2009). Toward reconstructing the evolution of advanced moths and butterflies (Lepidoptera: Ditrysia): an initial molecular study. BMC evolutionary biology 9.

Reynolds, S. E. and Bellward, K. (1989). Water Balance in *Manduca Sexta* Caterpillars: Water Recycling From the Rectum. Journal of Experimental Biology 141, 33.

Ritcher, P. O. (1958). Biology of Scarabaeidae. Annual Review of Entomology 3, 311–334.

Rossiter, M. C., Schultz, J. C. and Baldwin, I. T. (1988). Relationships among Defoliation, Red Oak Phenolics, and Gypsy Moth Growth and Reproduction. Ecology 69, 267–277.

Saini, R. S. (1964). Histology and physiology of the cryptonephridial system of insects. Transactions of the Royal Entomological Society of London 116, 347–392.

Scaccini, D. (2016). First record of oviposition scars in two European Platycerus species: *P. caprea* (De Geer, 1774) and *P. caraboides* (Linnaeus, 1758) (Coleoptera: Lucanidae). Bulletin de la Société royale belge d’Entomologie 152, 142–151.

Scaccini, D. (2022). Habitat and microhabitat suitability for Italian *Platycerus* species (Coleoptera: Lucanidae): elevation, slope aspect and deadwood features. Scandinavian Journal of Forest Research 37, 172–181.

Scaccini, D., Hendriks, P. and Fremlin, M. (2023). Morphometric traits of *Dorcus parallelipipedus* (Coleoptera: Lucanidae) larvae and adults: can differences between populations be ruled out? Annales de la Société entomologique de France 59, 249–262.

Schäfer, R. (1954). Zur Kenntnis der Anatomie, Physiologie und Ökologie des Brachkäfers, *Rhizotrogus aestivus Oliv*. (Col. Lam.). Zeitschrift für Angewandte Entomologie 35, 381–424.

Sheehan, C. M., Crawford, A. M. and Wigley, P. J. (1982). Anatomy and histology of the alimentary canal of the black beetle, *Heteronychus arator*. New Zealand Journal of Zoology 9, 381–385.

Shelomi, M., Lin, S. S. and Liu, L. Y. (2019). Transcriptome and microbiome of coconut rhinoceros beetle (*Oryctes rhinoceros*) larvae. BMC genomics 20.

Shields, V. D. C., Heinbockel, T. and Kristensen, N. P. (2006). Lepidoptera, Moths and Butterflies, Volume 2: Morphology, Physiology, and Development Handbook of Zoology: a Natural History of the Phyla of the Animal Kingdom Volume Iv Arthropoda: Insecta, Part 36. Annals of the Entomological Society of America 99, 988-989.

Sinha, R. N. (1958). The Alimentary Canal of the Adult of *Tribolium Castaneum* Herbst (Coleoptera, Tenebrionidae). Journal of the Kansas Entomological Society 31, 118–125.

Sirodot, M. S. (1858). Recherches sur les sécrétions chez les insects. Annales des sciences naturelles 10, 141–189.

Snodgrass, R. E. (1994). Principles of Insect Morphology. Ithaca, NY: Cornell University Press.

Soo Hoo, C. F. and Dudzinski, A. (1967). Digestion by the larva of the pruinose scarab, *Sericesthis geminata*. Entomologia Experimentalis et Applicata 10, 7–15.

Southwood, T. R. E. (1972). The insect/plant relationship - an evolutionary perspective. Symposia of the Royal Entomological Society of London 6, 3–30.

Stanbrook, R., Harris, E., Jones, M. and Wheater, C. P. (2021). The Effect of Dung Beetle Size on Soil Nutrient Mobilization in an Afrotropical Forest. Insects 12.

Stokland, J. (2012). Wood decomposition. In Biodiversity in Dead Wood (ed. J. Stokland, J. Siitonen & B. Jonsson), pp. 10–28. Cambridge: Cambridge University Press.

Strong, D. R., Lawton, J. H. and Southwood, R. (1984). 2 - The evolution of phytophagous insects. In Insects on plants : community patterns and mechanisms, pp. 15–45: Blackwell Scientific Publications.

Swingle, M. (1930). Anatomy and Physiology of the Digestive Tract of the Japanese Beetle. Journal of Agricultural Research 41, 181.

Sánchez, A., Micó, E., Galante, E. and Juárez, M. (2017). Chemical transformation of *Quercus* wood by *Cetonia* larvae (Coleoptera: Cetoniidae): An improvement of carbon and nitrogen available in saproxylic environments. European Journal of Soil Biology 78, 57–65.

Sánchez-Galván, I. R., Quinto, J., Micó, E., Galante, E. and Marcos-García, M. A. (2014). Facilitation among saproxylic insects inhabiting tree hollows in a Mediterranean forest: the case of cetonids (Coleoptera: Cetoniidae) and syrphids (Diptera: Syrphidae). Environmental Entomology 43, 336–343.

Takeshita, K. and Kikuchi, Y. (2020). Genomic Comparison of Insect Gut Symbionts from Divergent *Burkholderia* Subclades. Genes 11.

Takeshita, K., Matsuura, Y., Itoh, H., Navarro, R., Hori, T., Sone, T., Kamagata, Y., Mergaert, P. and Kikuchi, Y. (2015). *Burkholderia* of Plant-Beneficial Group are Symbiotically Associated with Bordered Plant Bugs (Heteroptera: Pyrrhocoroidea: Largidae). Microbes and environments 30, 321–329.

Tan, M., Wu, H., Yan, S. and Jiang, D. (2022). Evaluating the Toxic Effects of Tannic Acid Treatment on *Hyphantria cunea* Larvae. Insects 13.

Tanahashi, M. and Fremlin, M. (2013). The mystery of the lesser stag beetle Dorcus parallelipipedus (L.) (Coleoptera: Lucanidae) mycangium yeasts. *Bulletin of the Amateur Entomologists’* Society 72, 146–152.

Tanahashi, M. and Hawes, C. J. (2016). The presence of a mycangium in European *Sinodendron cylindricum* (Coleoptera: Lucanidae) and the associated yeast symbionts. Journal of insect science 16.

Tanahashi, M., Ikeda, H. and Kubota, K. (2018). Elementary budget of stag beetle larvae associated with selective utilization of nitrogen in decaying wood. The Science of Nature 105, 1–14.

Tanahashi, M., Kim, J. K., Watanabe, K., Fukatsu, T. and Kubota, K. (2017). Specificity and genetic diversity of xylose-fermenting Scheffersomyces yeasts associated with small blue stag beetles of the genus Platycerus in East Asia. Mycologia 109, 630–642.

Tanahashi, M. and Kubota, K. (2013). Utilization of the nutrients in the soluble and insoluble fractions of fungal mycelium by larvae of the stag beetle, *Dorcus rectus* (Coleoptera: Lucanidae). European Journal of Entomology 110, 611–615.

Tanahashi, M., Matsushita, N. and Togashi, K. (2009). Are stag beetles fungivorous? Journal of insect physiology 55, 983–988.

Tupy, J. H. and Machin, J. (1985). Transport characteristics of the isolated rectal complex of the mealworm *Tenebrio molitor*. Canadian Journal of Zoology 63, 1897–1903.

Ullman, D. E., Westcot, D. M., Hunter, W. B. and Mau, R. F. L. (1989). Internal anatomy and morphology of *Frankliniella occidentalis* (Pergande) (Thysanoptera: Thripidae) with special reference to interactions between thrips and tomato spotted wilt virus. International Journal of Insect Morphology and Embryology 18, 289–310.

Ulyshen, M. D. (2018). Saproxylic insects–diversity, ecology and conservation (1 edn): Springer Cham.

Vargas-Asensio, G., Pinto-Tomas, A., Rivera, B., Hernandez, M., Hernandez, C., Soto-Montero, S., Murillo, C., Sherman, D. H. and Tamayo-Castillo, G. (2014). Uncovering the cultivable microbial diversity of costa rican beetles and its ability to break down plant cell wall components. PloS one 9.

Verma, P. S. (1969). The alimentary canal and associated organs of two saprophagous beetle, Onthophagus catta Fabr. and Aphodius moestus Fabr. (Coleoptera: Scarabaeidae). Bulletin of entomology 10, 4–11.

Wada, N., Sunairi, M., Anzai, H., Iwata, R., Yamane, A. and Nakajima, M. (2014). Glycolytic Activities in the Larval Digestive Tract of *Trypoxylus dichotomus* (Coleoptera: Scarabaeidae). Insects 5, 351–363.

Wang, K., Gao, P., Geng, L., Liu, C., Zhang, J. and Shu, C. (2022). Lignocellulose degradation in *Protaetia brevitarsis* larvae digestive tract: refining on a tightly designed microbial fermentation production line. Microbiome 10.

Weihrauch, D. (2006). Active ammonia absorption in the midgut of the Tobacco hornworm *Manduca sexta* L.: transport studies and mRNA expression analysis of a Rhesus-like ammonia transporter. Insect biochemistry and molecular biology 36, 808–821.

Werner, E. (1926). Die ernährung der larve von potosia cuprea fbr. (Cetonia floricola hbst.). Zeitschrift für Morphologie und Ökologie der Tiere 6, 150–206.

Wessing, A. and Eichelberg, D. (1978). Malpighian tubules, rectal papillae and excretion. Genetics and Biology of Drosophila 2c, 1–42.

Wigglesworth, V. B. (1932). Memoirs: On the Function of the so-called ‘Rectal Glands’ of Insects. Quarterly Journal of Microscopical Science s2-75, 131–150.

Wigglesworth, V. B. (1974). The principles of insect physiology (Seventh edition.. edn). London: London: Chapman and Hall.

Yamada, A., Inoue, T., Noda, S., Hongoh, Y. and Ohkuma, M. (2007). Evolutionary trend of phylogenetic diversity of nitrogen fixation genes in the gut community of wood-feeding termites. Molecular ecology 16, 3768–3777.

Zhang, H. and Brune, A. (2004). Characterization and partial purification of proteinases from the highly alkaline midgut of the humivorous larvae of *Pachnoda ephippiata* (Coleoptera: Scarabaeidae). Soil Biology and Biochemistry 36, 435–442.

Zhang, L., Zhang, S., Ye, G. and Qin, X. (2020). Seasonal variation and ecological importance of tannin and nutrient concentrations in *Casuarina equisetifolia* branchlets and fine roots. Journal of Forestry Research 31, 1499–1508.

Šípek, P., Fabrizi, S., Eberle, J. and Ahrens, D. (2016). A molecular phylogeny of rose chafers (Coleoptera: Scarabaeidae: Cetoniinae) reveals a complex and concerted morphological evolution related to their flight mode. Molecular phylogenetics and evolution 101, 163–175.

